# Clinical risk factors of intensive care unit acquired weakness predicted by human muscle microtissue response to humoral factors

**DOI:** 10.1101/2025.10.23.684128

**Authors:** Heta Lad, Judy Correa, Kenneth Wu, Rebecca A. Gladdy, Margaret S. Herridge, Claudia C. dos Santos, Sunita Mathur, Jane Batt, Penney M. Gilbert

**Affiliations:** Institute of Biomedical Engineering, University of Toronto, Toronto, Canada; Donnelly Centre, University of Toronto, Toronto, Canada; Keenan Research Centre for Biomedical Science, St. Michael’s Hospital Unity Health Toronto, Toronto, Canada; Department of Medicine, Temerty Faculty of Medicine, University of Toronto, Toronto, ON, Canada; Lunenfeld-Tanenbaum Research Institute, Mount Sinai Hospital, Sinai Health, Toronto, Canada; Institute of Medical Science, University of Toronto, Toronto, Canada; Department of Surgical Oncology, Princess Margaret Cancer Centre/Mount Sinai Hospital, Sinai Health, Toronto, Canada; Department of Laboratory Medicine and Pathobiology, University of Toronto, Toronto, Canada; Department of Surgery, Mount Sinai Hospital, Sinai Health, Toronto, Canada; Division of Critical Care Medicine, University Health Network, Toronto, ON Canada; School of Rehabilitation Therapy, Queens University, Kingston, ON, Canada; Department of Cell and Systems Biology, University of Toronto, Toronto, Canada

## Abstract

ICUAW is an acquired phenomenon in the critically ill patient that is characterized by severe muscle weakness and atrophy. It results from both myopathic and neuropathic injury and is associated with heightened patient morbidity and mortality. Critical illness survivors exhibit variable functional outcomes following ICUAW, ranging from full recovery to persistent weakness with a significant negative impact on quality of life. Despite clinical importance, the mechanistic understanding of ICUAW remains incomplete, in part due to a paucity of tractable, species-specific experimental models. Previous studies have suggested a potential causative role of bloodborne factors, though this remains a matter of debate. To address these technological and knowledge gaps, we leveraged a 3-D human skeletal muscle microtissue (hMMT) culture platform to investigate the ability of humoral factors to directly impact pathogenesis of the myopathic component of ICUAW; critical illness myopathy (CIM). Blood serum collected from ICU patients within 72 hours of ICU admission was used to treat hMMTs for 6 days starting at a time-point when the myotubes were multinucleated, striated, and capable of generating force. hMMTs treated with serum from patients that survived the ICU stay exhibited several hallmarks of CIM including; significant reductions in myotube diameter (atrophy) and striation density, alongside functional declines in calcium release, peak force, and force kinetics. The *in vitro* profiles of ICU serum treated hMMTs displayed significant associations with known ICUAW and CIM clinical risk factors (e.g. age, length of ICU stay) such that individual patient risk factors could be predicted from serum-induced hMMT responses. An evaluation of hMMT responses enabled delineation of ICU non-survivors from healthy control and ICU survivors. This study offers an enabling technology for studies of CIM biology in the context of human cells while also delivering compelling evidence that humoral factors directly influence ICU-associated skeletal muscle pathogenesis. Together, these advances may inform strategies for preventing or treating ICUAW.

## 2. Introduction

Intensive care unit acquired weakness (ICUAW) develops in critically ill individuals due to varied combinations of both skeletal muscle and peripheral nerve dysfunction and injury, and referred to as critical illness myopathy (CIM) and critical illness polyneuropathy, respectively. ICUAW is characterized by profound weakness and atrophy of skeletal muscles with no obvious reason for weakness other than the ICU stay itself ^1–6^. CIM is a common contributor to ICUAW, with reported incidence averaging 40 % ^7–9^, and is associated with increased morbidity and mortality ^10,11^. Amongst ICU survivors, functional outcomes are heterogeneous; while some individuals recover fully to pre-ICU status, others will experience persistent weakness that may significantly reduce their quality of life ^10–13^. Various risk factors have been identified for the development of ICUAW. These include pre-existing health conditions, older age, and co-dependent factors including duration of mechanical ventilation (MV), length of ICU stay, and severity of acute illness^8,14^. Any exposure to systemic corticosteroids, sepsis, and multiorgan failure have also been implicated as potential contributors to ICUAW ^8,14^.

During the ICU stay, patients can lose 15 % of muscle mass within a single week, potentially leading to long lasting functional disability^15,16^. While muscle inactivity and unloading related to prolonged sedation, narcotic and paralytic use, and therapies administered in the ICU (e.g. corticosteroids) may contribute, they are not lone causative factors of ICUAW associated atrophy, the severity and hallmarks of which extend beyond these factors. Hallmarks of CIM include^17,18^ the preferential loss of thick filament protein myosin (MyHC), reductions in the MyHC/Actin ratio ^5,17–19^, and the loss of cross–striation patterning indicative of sarcomere disorganization ^20^. At a functional level, CIM has been characterized by non-excitable muscle membranes ^21^ and poor excitation-contraction coupling. These deficits manifest as a loss of specific force ^17,18^ and dysregulated calcium (Ca^2+^) release during force generation, such that an overabundance of Ca^2+^ is required to recruit the cross-bridges needed to activate contractile proteins and induce force generation ^18^. Despite this knowledge, there remains no early diagnosis or treatment for ICUAW as the underlying mechanisms driving the condition are not well understood. Thus, a pressing demand for robust experimental models to investigate the biology of the muscle and peripheral nerve injury underpinning ICUAW persists.

Porcine and rat ventilation models have been the primary modality for ICU basic research by simulating various CIM triggers and interventions such as prolonged MV, immobilization, sedation, corticosteroids, neuromuscular blocking agents and sepsis. Despite incompletely replicating the histological evidence of myopathy observed in human patients, the porcine model presents characteristic electrophysiological abnormalities associated with CIM, including significant decreases in compound muscle action potential (CMAP), and early muscle membrane dysfunction when ventilated for 5 days ^22–24^. Rats mechanically ventilated for > 5 days show a preferential loss of myosin in both slow and fast twitch muscles resembling CIM, in addition to the expected functional impairments ^25,26^. However, the practical application of these animal models is limited by the need for continual heavy sedation and paralysis during ventilation, as well as 24-hour observation and care, which incur significant cost and labor requirements.

Previous research suggests that ICUAW may be driven by humoral factors, i.e. factors transported in the blood ^27,28^. While humoral factors are unlikely to interact with muscle tissue in healthy individuals, there is an argument for its relevance in ICU patients where muscle edema is prevalent. In the ICU, several diagnostic techniques for CIM are reported as being difficult to perform due to complications owed to severe tissue edema^8^. Tissue edema occurs when there is an excessive amount of serous fluid trapped in or around cells and tissues of the body ^29^. The presence of edema implies “leaky” vasculature ^30^, suggesting that tissues may be exposed to more humoral factors than they would have prior to the onset of critical illness. There are a limited number of studies that investigate humoral factors in critically ill patients, which are largely sepsis-focused studies ^27,28^. Therefore, the specific role of humoral factors in ICUAW development is not well understood. With animal models, it is difficult to isolate the direct effects of humoral factors on skeletal muscle since indirect effects cannot be ruled out. Reducing complexity is a strength of culture-based studies. However, most *in vitro* muscle studies to date were conducted using 2D cultures of myotubes, where contractile apparatus maturation and muscle function assessments are fundamentally limited. Culture systems recapitulating ICUAW influences on human skeletal muscle tissue structure and function are presently lacking.

To address these gaps, we produced genetically identical arrays of human skeletal muscle microtissues (hMMTs) that we treated with serum collected from consenting critically ill patients, which we postulated would induce CIM-like phenotypes. hMMTs were fabricated within the skeletal muscle (Myo) microTissue Array deviCe To Investigate forCe (MyoTACTIC) 96-well culture plate, which supports non-invasive contractile force and calcium transient measurements of individual hMMTs over time ^31,32^. By implementing these real-time functional analyses together with end-point retrospective analyses (morphometric, protein studies), we proceeded to investigate the direct impact of ICU patient humoral factors (i.e. serum treatment) on engineered human skeletal muscle microtissues. Outcomes of hMMTs treated with each ICU patient serum sample were related to their corresponding demographic and ICU details to understand whether *in vitro* metrics were predictive of known ICUAW risk factors. Collectively, this work aimed to contribute to the long-standing debate on possible humoral origins of CIM, while also offering a first of its kind “CIM-in-a-dish” human cell-based culture system.

## 3. Materials and Methods

### 3.1 Study Design, Patient Recruitment, and Demographics

Individuals were recruited from the medical surgical and neurotrauma ICUs from St Michaels Hospital in Toronto, Ontario, Canada between October 2018 and May 2022. Informed consent was obtained from the patient or their substitute decision-maker. The study protocol was approved by the institutional research ethics board (REB 18-128). Subjects were eligible for study enrollment if they were 16 years of age or older, with acute onset of critical illness necessitating intubation and mechanical ventilation for an anticipated duration of 4 days or longer, and could be recruited within 72 hours of ICU admission. Subjects were excluded from enrollment if they presented with serious CNS injury with localized limb weakness (e.g. cerebrovascular accident, subarachnoid hemorrhage), a pre-existing formal diagnosis of neuromuscular disease, recent (prior 6 months) or current extremity trauma or surgery preventing movement and/or weight bearing (e.g. fracture, extensive soft tissue injury, joint replacement), were non-ambulatory before critical illness (either at home or in hospital), had known HIV, hepatitis B or C, or SARS COVID-19 infection (due to biosafety restrictions for serum experimentation in cell culture), anticipated death or withdrawal of care within 48 hours, or lack of a substitute decision maker if the patient was unable to consent. For hMMT investigation, blood serum was collected within 72 hours of ICU admission (i.e. admission serum).

892 patients were screened and 100 were eligible for study participation. Of these, 44 patients were consented to study participation, 49 patients refused, and 7 were repatriated or extubated prior to approach for consent. Of the 44 consented patients 1 died and consent was withdrawn for 1 prior to first blood draw, 1 was diagnosed post ICU admission with SARS-COV-2 infection necessitating sample exclusion, sera samples from 3 who died in the ICU could not be assayed due to technical reasons, and 2 patients had insufficient sample material. Consent was withdrawn, the patient was transferred to another hospital or palliative care or left the ICU against medical advice and thus was lost to follow-up for another 14 study participants. Of the remaining 22 patients, 5 died in the ICU, 1 was discharged home from the ICU, and 16 were discharged to the ward. Serum samples obtained from 22 patients at the time of ICU admission with known disposition were available for testing. Patient baseline demographics, clinical metrics, duration of ICU stay and treatment received are presented in Table 1. Serum collected from consenting healthy individuals was used for comparison purposes (Table 1). Cell culture experimentalist was blinded to patient data until all in vitro studies were completed.

**Table 1.**
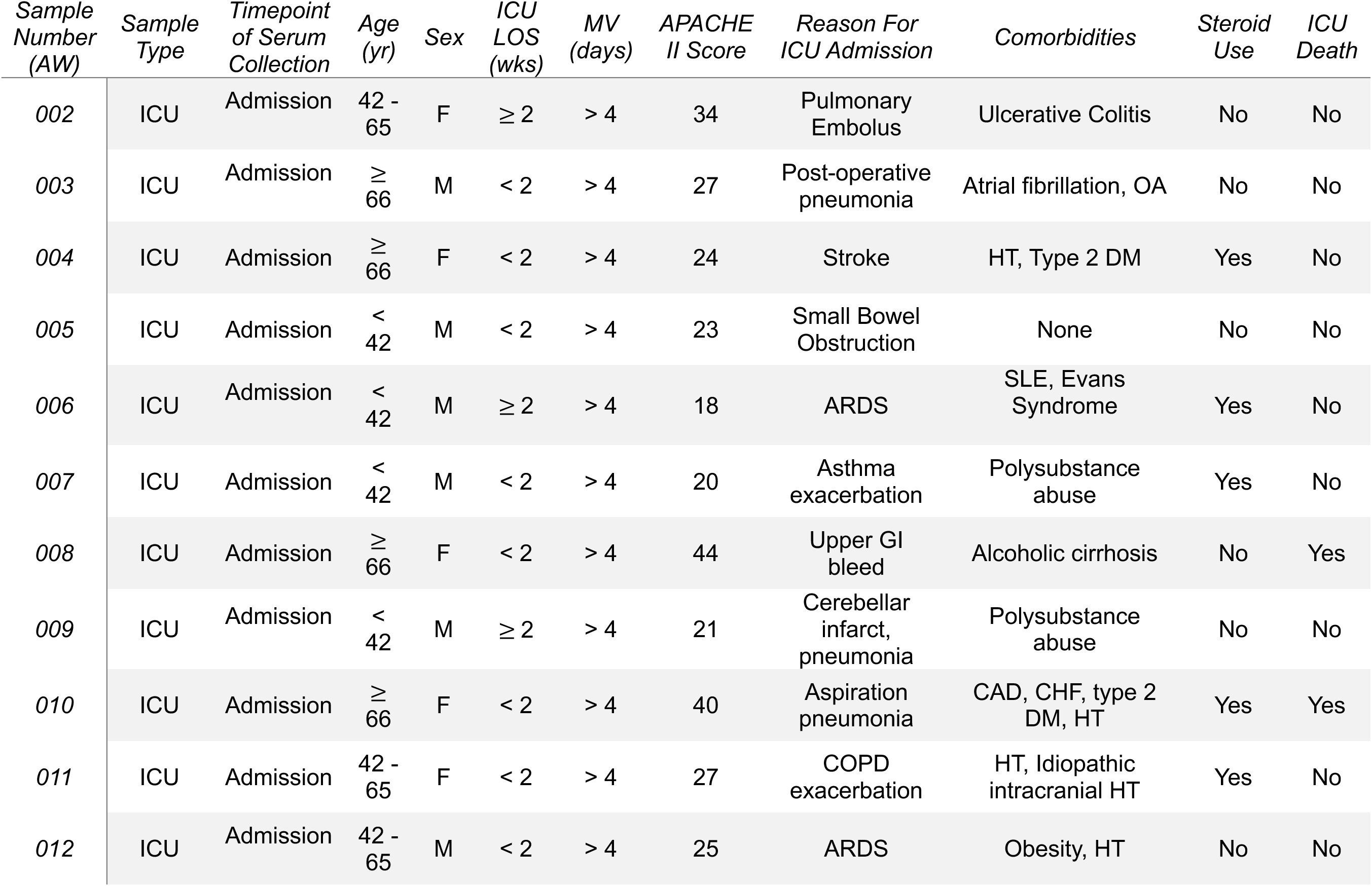
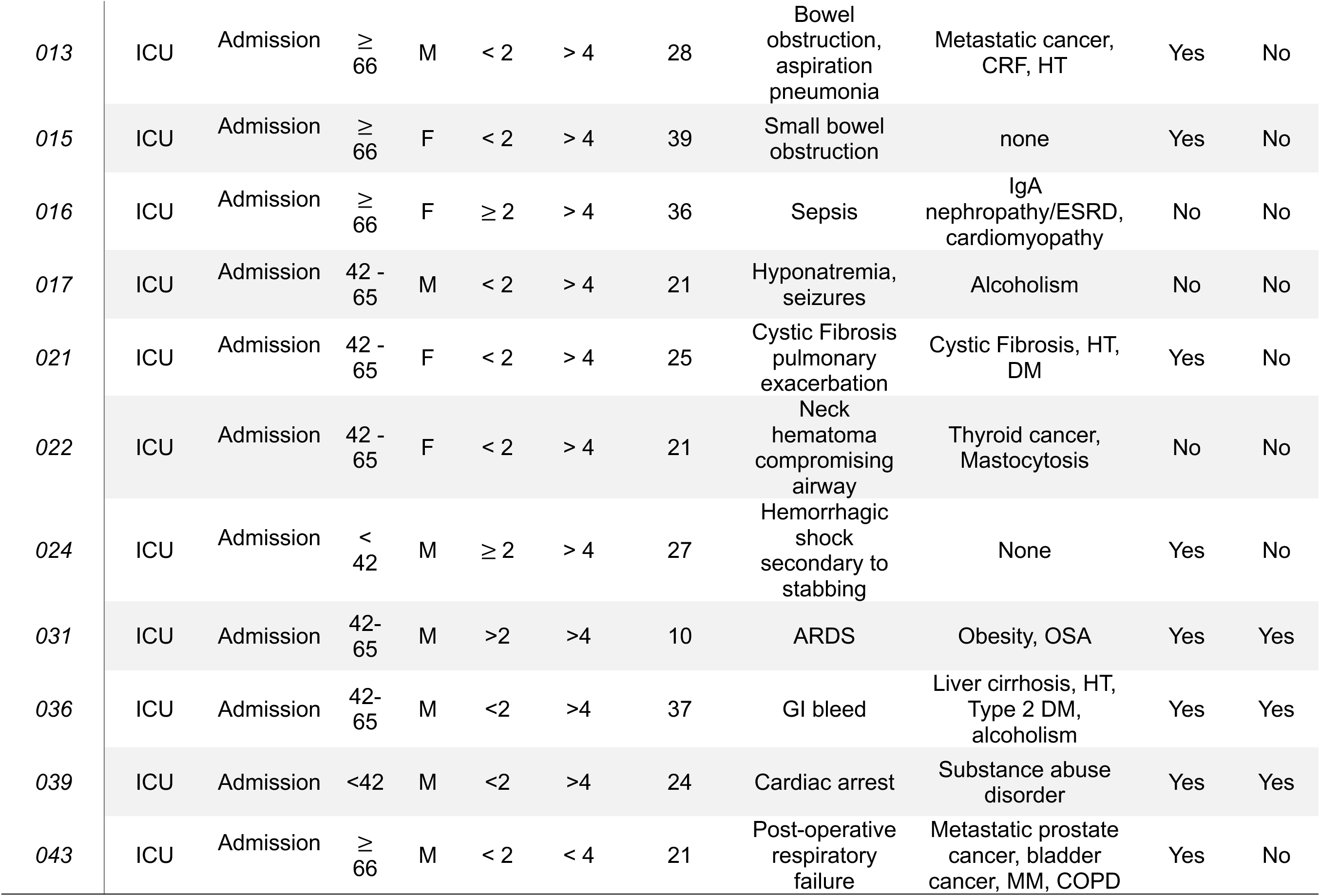
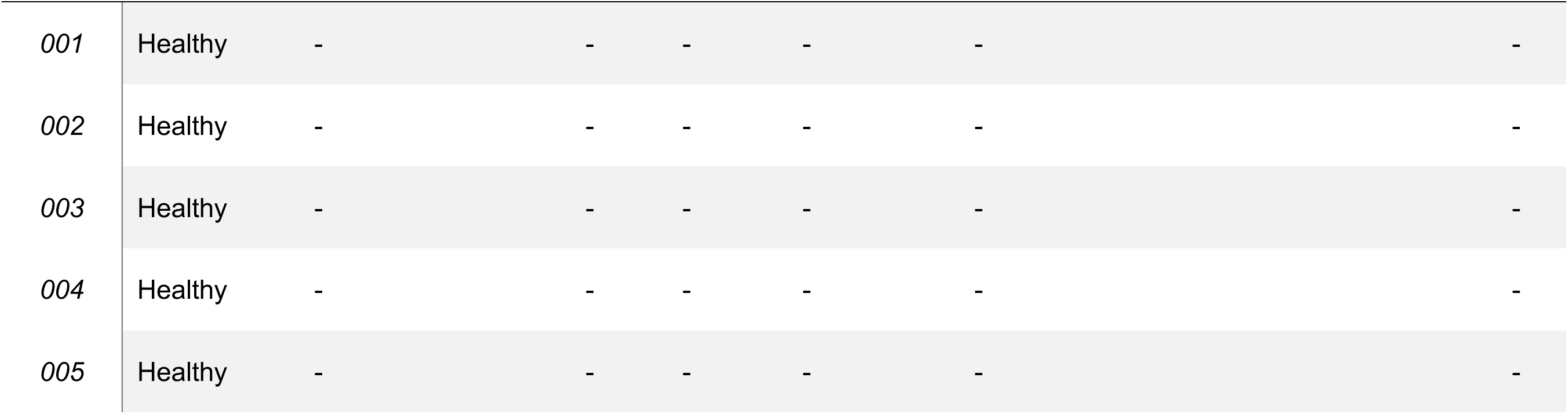
Patient Demographics.

### 3.2 Rectus Femoris Ultrasound, Medical Research Council Sum Score and Functional Independence Measure Score

B-mode ultrasound (General Electric Logiq *e*; GE Medical Systems, Milwaukee, WI) of the quadriceps muscle was performed by a trained operator to obtain muscle thickness and cross-sectional area of the rectus femoris (RF) muscle at 50% of thigh length (from ASIS to superior pole of patella). The ultrasound frequency was set at 12 MHz, gain (58 – 72 dB) and depth (4-6 cm) settings were optimized by the operator to obtain a clear image of the muscle. Captured images were exported from the ultrasound system in DICOM format to Osirix MD software (https://www.osirix-viewer.com/osirix/osirix-md/), and manually outlined to obtain measures of muscle thickness or cross-sectional area as previously described ^33,34^. The average measurement from three images was used for statistical analysis. Imaging was performed at ICU admission, every 4 to 5 days during ICU stay, pending patient clinical stability and operator availability.

Global muscle strength was manually assessed with Medical Research Council (MRC) testing by a trained assessor (physical therapist) and the MRC Sum Score (MRCSS) was calculated ^35^. MRCSS ranges from 0 to 60, with 60 indicating “normal” muscle strength (age and sex appropriate). MRCSS testing was completed in the ICU on study participants when awake and fully cooperative. Physical functional capacity was assessed by determining the motor component of the Functional Independence Measure (FIM) at 7 (+/- 2) days post ICU discharge in critical illness survivors.

### 3.3 Human Immortalized Myoblast Culture

The AB1167 human immortalized myoblast cell line used in this study was established by MyoLine at the Institut de Myologie (Paris, France) ^36^. More details regarding the maintenance of this culture line can be found in our previous reports ^32,37^. In brief, immortalized human myoblasts were seeded into 10 cm tissue culture plates (3 x 10^5^ cells per dish) with myoblast growth media containing 84 % Skeletal Muscle Cell Basal Medium with Skeletal Muscle Cell Growth Medium Supplement Mix (Sigma Aldrich, Cat. #C-23260), supplemented with 15 % fetal bovine serum (FBS; Gibco, Cat. #12483020), and 1 % penicillin/ streptomycin (Gibco, Cat. #15140122). The full myoblast growth media was exchanged with fresh media every 2 days during the culture period. Immortalized myoblast cultures were maintained until 75 % confluency, after which the cells were either sub-cultured into new tissue culture dishes or used to generate 3D human skeletal muscle microtissues (hMMTs) as described below. Immortalized myoblasts underwent no more than 14 expansion passages before being seeded into hMMTs.

### 3.4 MyoTACTIC Platform Fabrication

MyoTACTIC platform fabrication was completed using polydimethylsiloxane (PDMS; Sylgard^TM^ 184 silicone elastomer kit, Dow Corning; Midland, MI, USA). All plate features were cast in a single step from a reusable, negative polyurethane (PU; Smooth-Cast® 310 liquid plastic, Smooth-On; Macungie, PA, USA) mold. More detailed methods on how to generate a reusable PU mold and fabricate our PDMS plate have been previously reported by our group and all supporting information can be found on our GitHub (https://github.com/gilbertlabcode) ^31,32^. PDMS MyoTACTIC plates were first cut into smaller functional units and prepared for tissue seeding by sonicating MyoTACTIC portions for 25 minutes in isopropanol (Cat. 8600, Caledon; Georgetown, Ontario, Canada), after which they were rinsed twice in ddH2O and set to evaporate in a curing oven for 15 minutes. MyoTACTIC portions were autoclaved for sterilization before use.

### 3.5 Human Skeletal Muscle Microtissue (hMMT) Fabrication and Treatment

hMMTs were fabricated based on our previously described methods ^31,32^. In brief, immortalized myoblasts were gently dissociated from their culture plates using 0.25 % Trypsin-EDTA (Thermo Fisher Scientific, 25200072) and then resuspended in a protein-rich hydrogel mixture containing 4 mg/mL fibrinogen (40 % v/v; Sigma-Aldrich, F8630) and Geltrex (20 % v/v; Thermo Fisher Scientific, A1413202) in DMEM (40 % v/v; Gibco, 11995-065). Fibrinogen from bovine plasma was reconstituted in 0.9 % sterile saline solution (house brand) at 37 °C for 3 mins and subsequently filtered using a 0.2 μm sterile filter before use. The cell suspension arrived at a final density of 1.5 × 10^5^ cells per tissue. Thrombin from human plasma was added to the cell suspension at a concentration of 0.2 units of thrombin per mg of fibrinogen, then the mixture was quickly pipetted into the oval pool of each MyoTACTIC well ensuring the cell suspension was spread evenly across the surface of the oval pool and surrounded the PDMS posts. The seeded MyoTACTIC wells were placed in the 37 °C incubator for 5 minutes to expedite fibrin gel formation. After hydrogel formation, each hMMT was supplemented with 200 μl of growth media (GM) containing Promocell skeletal muscle basal media (no supplement mixture) supplemented with 20 % FBS, 1 % Penicillin/Streptomycin, and 1.5 mg/ml 6-aminocaproic acid (3 % v/v; (Sigma-Aldrich, #A2504)). This time point is referred to as day -2. Two days later, and referred to as differentiation day 0, GM was removed and differentiation media (DM) containing 2 % horse serum (HS; Gibco, #16050-122), 2 mg/ml 6-aminocaproic acid (4 % v/v) and 10 μg/ml human recombinant insulin (Sigma Aldrich, Cat# 91077C) in DMEM was added. Half of the hMMT differentiation media was refreshed every other day until the experimental endpoint, day 12 of differentiation for all the hMMTs. All media compositions are highlighted in Table 2.

**Table 2.** Composition of media and solutions.

| Media/Solutions | Composition |
| --- | --- |
| Freezing Media | 90 % FBS* (Gibco, #12483020,), 10% Dimethyl Sulfoxide (Sigma-Aldrich, #D8418) |
| Wash Media | 89 % DMEM* (Gibco, #11995073), 10 % FBS, 1 % P/S* (Gibco, #15140122) |
| Growth Media | 84 % Skeletal Muscle Cell Medium Ready-to-Use (Promo-Cell via Sigma-Aldrich, #C-23260), 15 % FBS, 1% P/S |
| After Seeding Growth Media (hMMTs) | Skeletal Muscle Cell Basal Medium, 20 % FBS* 1.5 mg/mL ACA* (3 % v/v; Sigma-Aldrich, #A504), 1 % P/S |
| Differentiation Media (hMMTs) | 92 % DMEM*, 2 % HS* (Gibco, #16050-122), 2 mg/mL ACA* (4 % v/v), 10 µg/mL human recombinant insulin (Sigma-Aldrich, #91077C), 1 % P/S* |
| Fibrinogen Solution | 10 mg/mL fibrinogen (Sigma-Aldrich, #F8630) in 0.9% (wt/v) NaCl solution in ddH<sub>2</sub>O |
| Extracellular Matrix Solution | 40 % DMEM (v/v), 4 mg/mL fibrinogen (40 % v/v), 20 % Geltrex™ (v/v; Gibco, #A1413202) |
| Permeabilization and Blocking Solution (Immunostaining) | 10 % GS* (Gibco, #16210072), 0.3 % Triton X-100 (BioShop, #TRX777) in D-PBS |
| Calcium Handling Solution | Hanks' buffer with 20 mM Hepes (Cedarlane, #20011(AAT)), 0.04 % Pluronic Acid (w/v), 10 µM Calbryte™ 520 AM (Cedarlane, # 20653(AAT)) |
| Lysis Buffer 1 | 25 mM Tris (BioShop, TRS001.1) pH 7.2, 0.5% Triton X – 100, in ddH2O |
| Lysis Buffer 2 | 20 mM Tris pH 7.2, 100 mM potassium chloride (Sigma-Aldrich, P3911), 20% glycerol (BioShop, GLY002.1), 1 mM DTT (BioShop, DTT002.5), 50 mM spermidine (Sigma-Aldrich, S0266) in ddH2O |
| TBS-T | Tris-buffered saline with 1 % Tween 20 (BioShop, TWN510.500) |
| Blocking Solution (Western Blotting) | 5 % Skim milk (w/v; BioShop, SKI400) in TBS-T |
| 10x Running Buffer (1L) | 30.3 g Tris base, 144.1 g glycine (BioShop, GLN001), 50 ml Sodium dodecyl sulfate (20 % w/v; BioShop, SDS001), in ddH2O |
| 1x Running Buffer | 100 ml of 10 x running buffer in 900 ml of ddH2O |
| 10x Transfer Buffer (500 ml) | 15.14 g Tris base, 72 g glycine in ddH2O |
| 1x Transfer Buffer | 100 ml of 10 x transfer buffer, 200 ml methanol in 700 ml ddH2O |

\*\*Dulbecco's Modified Eagle's Medium (DMEM). \*6-Aminocaproic acid (ACA). \*Penicillin-Streptomycin (P/S). \*Fetal bovine serum (FBS). \*Goat Serum (GS).

For the healthy human serum (HuS) analysis, hMMTs were cultured in the standard differentiation media indicated above. This occurred until day 6 of differentiation. On day 6 of differentiation, the entire volume of DM was removed from each MyoTACTIC well and new DM containing 2 % of healthy HuS, instead of HS, was supplemented into the designated tissues. After that, all half media changes contained healthy HuS. For ICU patient studies, on day 6 of differentiation, the DM was completely removed. New DM containing 2 % admission serum (ICU – HuS) or 2 % healthy HuS, instead of HS was supplemented. After that, all half media changes were with DM containing ICU patient serum or the healthy HuS control. This allowed for 6 days of hMMT treatment with the ICU patient serum samples. With this protocol, we prioritize exposure at a timepoint when myotube myonuclear accretion has ceased (DM6) while maximizing sera exposure time (6 days) before hMMT integrity declines, which is observed starting in the third week of DM culture.

### 3.6 Tissue Compaction Analysis

Phase-contrast images of all hMMTs were acquired with the cellSens^TM^ software every other day of differentiation before the media was refreshed. A single image of each hMMT was taken at 4X magnification, ensuring that the entire hMMT (post to post) fit in the field of view. Images were analyzed using the NIH FIJI/ImageJ software ^38^. The effect of increasing serum dosages on hMMT compaction was evaluated by measuring hMMT width as an indication of tissue self-organization and remodeling over the differentiation period. For each hMMT, the width of the microtissue was measured in three regions, one on either end of the hMMT near the posts and one measurement down the center of the microtissue. Measurements were averaged, and then pixel values were converted into microns (295 pixels = 500 μm). This was done for days 4, 6, 10, and 12 of differentiation, representative of hMMT width throughout differentiation, before and after healthy HuS was introduced.

### 3.7 Immunostaining, Myotube Width, Coefficient of Variance and Striation Analysis

To visualize myotubes within 3D hMMTs we performed immunostaining. Antibody vendor information can be found in Table 3. On day 12 of differentiation, following aspiration of culture media, hMMTs were washed with D-PBS for 3 x 2 mins, and then fixed in 4 % paraformaldehyde (PFA) in D-PBS at room temperature for 15 minutes. Following D-PBS washes, hMMTs were simultaneously blocked and permeabilized using a solution of 0.3 % Triton-X 100 (BioShop, TRX777.500) and 10 % goat serum (Gibco, 16210072) in PBS. Blocking was done for 30 - 45 minutes at room temperature. Subsequently, hMMTs were incubated with a mouse anti-sarcomeric α-actinin antibody (SAA, 1:800; Sigma, A7811) diluted in the blocking solution overnight at 4 °C. After washing with D-PBS three times for 10 mins each, samples were then incubated for 45 mins at RT with goat anti-mouse IgG conjugated with Alexa-Fluor 488 antibody (1:500; Invitrogen, A-21131) and Hoechst 33342 for counterstaining nuclei (1:1000; Cell Signalling, 4082S) diluted in the blocking solution. Thereafter, hMMTs were washed 3 times for 10 minutes in D-PBS to remove any excess secondary antibody, after which hMMTs for ready for fluorescence imaging.

**Table 3.** List of antibodies.

| Antibody | Host | Dilution | Use | Manufacturer |
| --- | --- | --- | --- | --- |
| anti-sarcomeric α-actinin | Mouse | 1:500 | Immunostaining,<br>western blot | Sigma-Aldrich,<br>#A7811 |
| Hoechst 3342 | - | 1:1000 | Immunostaining | Thermo Fisher<br>Scientific, #H3570 |
| Goat anti Rabbit IgG<br>(H+L) Secondary<br>Antibody, Alexa Fluor<br>488 | Goat | 1:800 | Immunostaining | Invitrogen, #A-11008 |
| pAMPK-α (Thr172)<br>(40H9) | Rabbit | 1:1000 | Western Blot | Cell Signalling,<br>#2535 |
| AMPK-α | Rabbit | 1:1000 | Western Blot | Cell Signalling,<br>#2532 |
| PGC-1α | Rabbit | 1:500 | Western Blot | Sigma Aldrich,<br>#AB3242 |
| pan-myosin heavy chain | Mouse | 0.5<br>μg/ml | Western blot | DSHB, MF20 |
| actin | Mouse | 0.5<br>μg/ml | Western blot | DSHB, JLA20 |
| GAPDH | Mouse | 0.5<br>μg/ml | Western Blot | DSHB, GADPH-2G7 |
| anti-mouse IgG, HRP | Horse | 1:2000 | Western Blot | Cell Signalling,<br>#7076S |
| anti-rabbit IgG, HRP | Goat | 1:2000 | Western Blot | Cell Signalling,<br>#7074 |

To examine myotube formation in hMMTs, 40x confocal fluorescent images were acquired using Fluoview-10 imaging software (FV-10; Olympus) and an IX83 inverted microscope (Olympus). Five confocal images were captured per hMMT at five, non-overlapping locations across each hMMT. These locations were kept consistent for each hMMT analyzed and determined before imaging. Confocal image stacks of hMMT immunofluorescence were analyzed using the NIH ImageJ software. Non-damaged myotubes, visible across the length of a 40x image, were considered for width analysis if they had three or more nuclei. The diameter of each myotube was taken at 3 places and averaged to obtain the value of a single myotube. A minimum of 5 myotubes were analyzed per image where possible. The myotube diameter was averaged for each image stack. This was repeated for all 5 images acquired for a single hMMT. Next, the average myotube diameter was averaged across all 5 images to obtain the mean myotube diameter for an entire hMMT. This was repeated for all hMMTs. The coefficient of variance for each myotube was determined by dividing the standard deviation for each myotube, from its respective average myotube width. A smaller value is indicative of a smaller variation, while a larger value is indicative of a larger variation. This was repeated for all myotubes analyzed in each hMMT confocal image.

Each myotube analyzed for myotube diameter was also investigated for whether striated patterning was present or absent. In some cases, myotubes are not striated along their full length. Therefore, if a myotube was > 50 % striated, it was characterized as striated. The following formula was used to determine the percentage of striated myotubes: % *of striated myotubes* = (# *of striated myotubes* / *total* # *of myotubes*) *x* 100

### 3.8 hMMT Myonuclear Fusion FusionIndex

hMMT nuclear fusion index was analyzed by assessing 40x magnification confocal images from SAA immunostained and Hoechst counterstained hMMTs on day 12 of differentiation. For this, we used MyoCount, an open-source programs that runs analysis using MATLAB runtime ^39^. The tool identifies myotubes containing at least *n* nuclei (default setting *n=3*). It reports the following metrics for an image: the total number of nuclei, the total number of nuclei within myotubes, and the total number of nuclei within myotubes containing at least *n* nuclei. A total of 2 images per hMMT were run. Default metrics were optimized to fit our images and the following metrics were implemented:

- MinCircleRad 10
- MaxCircleRad 35
- TubeThresh 0.8
- NucFillSize 15
- SmallestNucleusPixelCount 80

ImageJ was used to facilitate the quantification of SAA coverage in z-stack confocal images of each hMMT projected over maximum intensity. The SAA channel was split from the Hoechst channel and used for the remainder of this analysis. First, the SAA signal was pseudo-coloured as red, then the image was converted to RGB format. To remove background noise and increase the red pseudo-coloured SAA signal, the image was manually thresholded. Subsequently, a modified ImageJ Macro originally developed by McColl *et al* ^40^, and reported here^37^, was run to produce the inverse area coverage of the SAA signal. This value was subtracted from 100 to obtain the percentage of SAA coverage for each image. Therefore, SAA coverage was averaged across all images for each hMMT.

### 3.9 hMMT Passive Force Analysis

On differentiation days 6 and 12, phase-contrast images were taken of each hMMT at 4x magnification, ensuring focus was on the myoTACTIC posts. It is critical to keep this focal plane and position consistent for each image acquisition and this was done by focusing on a single select post for each hMMT each time an image was acquired. Images were then analyzed using the NIH ImageJ software. A single measurement was taken from one post to the other, across the length of the hMMT. The location of this post to post measurement was kept consistent for differentiation days 6 and 12 for each hMMT. Then, pixel values corresponding to the distance across the length of the tissue were converted into microns (295 pixels = 500 μm). To determine the change in passive force, the measurement taken on day 6 (in μm) was subtracted from the day 12 value for each tissue and then converted into absolute passive force (μN). Positive values were indicative of a larger intracellular tension, while negative values were indicative of a decrease in intracellular tension.

### 3.10 hMMT Electrical Stimulation Optimization and Contractile Force Measurement

Electrical stimulation (E-Stim) parameters of voltage, frequency, and duty cycle were optimized to determine AB1167 hMMT peak twitch (0.5 Hz, 5 Vpp, 3 % / 80 ms duty cycle) and tetanus force (60 Hz, 5 VPP, 4 % / 5 ms duty cycle) ^41^. In-depth methodological information on setup and analysis can be found in ^31,32^. In brief, on differentiation day 11 for ICU patient studies, and day 12 for all other force studies, two regular bevel needles, acting as electrodes, were inserted behind each post in a single MyoTACTIC well. The “electrodes” were then connected to a commercial pulse generator using tinned copper wire, and alligator clamps. Movies of post-deflection were captured using an Apple iPhone SE during electrical stimulation of hMMTs at 10X magnification using an Olympus confocal IX83 microscope and a LabCam™ iPhone microscope mount. For contractile force measurements, hMMTs were induced to contract 6 times with the optimized stimulation parameters. The contractile force of the first contraction is often irregular in comparison to subsequent contractions, and thus was excluded from the analysis. Peak force values were averaged over contractions 3-5 to correspond to the quantification of Twitch Force. For stimulation at 60 Hz corresponding to tetanic force, we report the maximum absolute force as the absolute force generated during the second tetanic contraction. This pixel value was averaged and converted to obtain absolute contractile forces generated by the hMMTs (µN). For these experiments, four hMMTs from a single MyoTACTIC portion served as replicates. Quantification of E-Stim videos was completed using the semi-automated script found on our GitHub URL as reported here ^31,32^.

### 3.11 hMMT Induced Calcium Transient Analysis

On day 12 of differentiation, DM was removed from hMMTs and kept aside. hMMTs were washed 2 x 3 minutes with warmed PBS, then incubated for 1 hour in the 37 °C incubator with Calbryte^TM^ 520 AM (Cedarlane, # 20653(AAT)) at 10 uM in hanks buffered saline with 20 mM Hepes (Cedarlane, # 20011(AAT)) supplemented with 0.04 % pluronic acid (v/v). After incubation, the calcium indicator was removed and hMMTs were washed for 2 x 3 minutes with warmed PBS. hMMTs were then returned to their DM and allowed to warm for 15 mins in the 37 °C tissue culture incubator. Calcium transients were captured in response to E-Stim (see details above; twitch and tetanus) using an inverted microscope (Olympus IX83) with a 4x magnification lens. Consecutive time-lapse epi-fluorescent images were captured at a frame speed of ∼135 ms with a DP80 CCD camera (Olympus) outfitted with a fluorescein isothiocyanate (FITC) filter and Olympus cellSens^TM^ imaging software. Subsequently, the fluorescent signal in the series of time-lapse images was measured in a single region of interest (ROI) within the center of each hMMT using Fiji software (ImageJ, NIH), and relative fluctuations in the fluorescence signal were calculated and presented as ΔF/F_0_ = (F_immediate_ – F_baseline_)/(F_baseline_). The relative change in fluorescence signal for each hMMT in response to twitch and tetanus contractions was determined and plotted against time. Integrated density, a measure of total calcium released, was measured as the sum of the Mean Grey Value x Area for each frame during a 2-second tetanic stimulation period using Fiji software (ImageJ, NIH). For all investigations, each tissue was normalized to its minimum before comparison. Studies to validate that the calcium indicator was reporting Ca^2+^ released from the sarcoplasmic reticulum were conducted by treating hMMTs with 75 mM of dantrolene sodium (Sigma, #251680).

### 3.12 hMMT Digestion and Western Blotting Analysis

Western blotting was performed on hMMT microtissue samples collected on day 12 of differentiation, immediately downstream of calcium handling analysis. Lysate from a native human muscle biopsy was included as an antibody positive control. The collection and use of this human skeletal muscle tissue was reviewed and approved by the Mount Sinai Hospital Research Ethics Board (REB14-0109-E). The University of Toronto Office of Research Ethics also reviewed the approved this study and assigned administrative approval (Protocol# 30557). All procedures in this study were conducted in accordance with the guidelines and regulations of both Research Ethics Boards. Patient consent was obtained. The tissue digestion methods were adapted from Roberts *et al* ^42^. In brief, following removal of DM, hMMTs were washed 2 x 3 minutes with D-PBS and all the technical replicates of a single treatment were transferred into the same 1.5 ml Eppendorf tube. Using a micro-tube homogenizer system (VWR, 76529-604), outfitted with a disposable pellet pestle, for 1.5 mL microtubes (Fisher Scientific, FS7495211590), hMMTs were homogenized for ∼ 15 seconds without lysis buffer to first break down the tissue. Next, hMMTs were homogenized for ∼ 15-30 seconds in buffer 1 containing 25 mM Tris (BioShop, TRS001.1) pH 7.2, 0.5% Triton X – 100. Following a spin at 1500 g X 12 minutes, the cytoplasmic fraction supernatant was transferred into a new Eppendorf tube and kept at -80 °C until use. The remaining pellets containing the myofibrillar fraction (MF) were washed in buffer 1 at 1500 g x 7 minutes. After centrifugation, the supernatant was discarded and the pellet was resuspended in buffer 2 containing 20 mM Tris pH 7.2, 100 mM potassium chloride (Sigma-Aldrich, P3911), 20% glycerol (BioShop, GLY002.1), 1 mM DTT (BioShop, DTT002.5), 50 mM spermidine (Sigma-Aldrich, S0266) using a new pellet pestle. Samples were spun down for 1 minute using a portable bench-top centrifuge and the supernatant containing the myofibrillar fraction (MF) was transferred to a new tube and stored at -80 °C until use. The total protein content for each sample was measured using the BCA assay kit (Thermo Fisher Scientific, 23227).

5 μg of total hMMT MF protein or 15 μg of cytoplasmic protein, alongside 0.5 μg of human positive control protein (MF or cytoplasmic) was separated on a Tris-Glycine SDS-PAGE gel (8 %) and then transferred into a nitrocellulose membrane (BioRad, 1704158) using a wet transfer system. Membranes were washed with ddH2O immediately after being transferred and then incubated in ponceau stain (BioShop, PON002.500) for 15 mins. After 2 x 3 minutes washes with ddH_2_0, blots were imaged using the BioRad ChemiDoc^TM^ MP imaging system to obtain total protein stain for use as loading controls. Membranes were then washed with Tris-buffered saline with 1 % Tween 20 (TBS-T; BioShop, TWN510.500) and then blocked with a blocking solution containing 5 % skimmed milk in TBS-T for 30 minutes at room temperature. Then, membranes were incubated with primary antibodies against skeletal muscle sarcomeric proteins including mouse anti-myosin heavy chain-pan (MF-20; DSHB, 0.5 µg/mL), mouse anti-sarcomeric alpha-actinin (1:2000; Sigma) and mouse anti-actin (JLA20; DSHB, 0.5 µg/mL), or cytoplasmic proteins PGC-1α (1:500, Sigma), pAMPK (1:1000, Cell Signalling), and AMPK (1:1000, Cell Signalling) diluted in the blocking solution overnight at 4 °C. Subsequently, membranes were washed with TBS-T (3 x 5 minutes) and incubated with horseradish peroxidase-conjugated secondary antibody, goat anti-mouse IgG (1:2000, Cell Signaling) or goat anti-rabbit IgG (1:2000, Cell Signalling) diluted in the blocking solution for 30 minutes at room temperature. Finally, membranes were washed with TBS-T (3 x 5 minutes) and developed using SuperSignal™ West Dura Chemiluminescent Substrate (Thermo Fisher Scientific, 0034075) and imaged with the BioRad ChemiDoc^TM^ MP imaging system. Blot quantification was done using the NIH ImageJ software. All buffer and antibody compositions are provided in Table 2.

### 3.13 hMMT Metric Clustering with Binary Clinical Data

All admission hMMT metrics were normalized to their respective healthy controls and compiled into a single Excel sheet. Using Excel, we applied conditional formatting, and new rules were generated. We formatted cells based on their values implementing a 3-colour scale. Using the ‘type’ number, we set a threshold. A value of 1 specifies equivalency to the healthy HuS control and will appear white. Values > 1 are orange (darker shades for higher values), and < 1 are blue (darker shades for lower values).

We set thresholds at 1.5 (darkest orange) and 0.5 (darkest blue). Each heatmap contains 2 clusters (left, admission; right: 7DPID). Supporting bar plots were generated in Jupyter Notebook (v. 6.5.4) implementing the ‘pandas’, ‘seaborn’, ‘matplotlib.pyplot’ and ‘scipy.stats’ libraries.

### 3.14 hMMT Statistical Analysis

Statistical analysis was performed using GraphPad Prism 9.0 (GraphPad Software; Boston, MA, USA) or computed in Jupyter notebook (v. 6.5.4). For data with multiple groups compared across one independent variable, a one-way or two-way ANOVA followed by Tukey’s multiple comparison test was utilized to compare the means of all groups to one another. When the effects between two variables were compared, a unpaired student t-test was used to determine significance. All values are expressed as mean ± standard error of the mean (SEM) using ‘n’. Significance was defined as p ≤ 0.05. Files containing all raw data have been provided.

Supporting bar graphs for heatmaps were generated in the Jupyter Notebook (v. 6.5.4). Baseline metrics from admission data were analyzed across age, age and length of stay (LOS), or steroid use, to assess differences using non-parametric tests due to the distributional properties of the data. The Kruskal-Wallis test was initially employed to evaluate overall differences among age groups, and age plus LOS for each metric. Subsequently, pairwise comparisons were conducted using Tukey’s Honestly Significant Difference (HSD) test to identify specific group differences. In studies where patients were grouped by steroid use status (Steroid vs. No Steroid) the Mann-Whitney U test was utilized for its robustness against non-normally distributed data and its ability to assess differences between independent groups. Statistical significance was determined at a significance level of 0.05 for all plots. Raw data points were plotted alongside median values to visualize the distribution of individual observations within each group. Python 3 and associated libraries were utilized for statistical computations and visualizations. Data preprocessing and statistical analyses were conducted using Pandas, Matplotlib, SciPy, and Statsmodels libraries.

## 4. Results

### 4.1 hMMTs differentiated with 2 % healthy HuS are structurally and functionally comparable to the established standard of 2 % HS

We first confirmed that replacing the 2 % horse serum (HS; established standard) in differentiation media (DM) with healthy human serum (HuS) had no gross impact on hMMT structure or function. For this, all hMMTs were differentiated in standard DM containing 2 % HS until day 6 of differentiation when myotubes have formed and are capable of generating force (Figure S1A). At this point, the standard DM was replaced with DM containing 2 % HuS. Half-media changes were performed every other day thereafter until the day 12 study endpoint (Figure S1A). No significant differences in tissue formation (S1B), compaction over time (S1C), gross morphology (S1D) or in the width of myotubes within hMMTs (S1E) were found upon comparing hMMTs cultured in DM containing 2 % HS to those with 2 % HuS.

We next confirmed that all of our control healthy HuS samples, individually or mixed together, did not significantly alter the structural and functional properties of hMMT myotubes as compared to the established standard (HS). hMMTs were first subject to morphometric analysis via immunostaining. Sarcomeric α–actinin (SAA) and Hoechst 33342 (nuclei) immunostaining at day 12 of differentiation revealed hMMTs with large, multinucleated, and striated myotubes comparable across all healthy HuS samples, individually or mixed, in comparison to HS (Figure S2A-C). Next, we analyzed passive force, which reflects the tension exerted on flexible rubber posts by tissue intracellular tension. Increased intracellular tension is expected since myotubes continue to mature during hMMT differentiation (Figure S2D), while compromised myotube integrity reduces tension and decreases passive force (Figure S2D). Quantifying the distance between the posts via phase contrast images revealed that all values remained positive (i.e. increase in tension), with no significant differences observed across the healthy HuS samples compared to HS (Figures S2E). Finally, we investigated the active contractile force of hMMTs in response to electrical stimulation (E-Stim). Quantification of post displacement revealed no significant difference in absolute twitch and tetanic forces in hMMTs across all healthy HuS tested independently or mixed together, as compared to HS (Figure S2F-G). These studies confirmed the structural and functional similarity of hMMTs generated with healthy HuS as compared with HS.

### 4.2 6-day treatment with admission serum from ICU survivors impairs hMMT structural integrity

We then proceeded to test the hypothesis that exposing hMMTs to serum collected from patients within the first 72 hours of admission to the ICU is sufficient to induce CIM-associated phenotypes. We initially focused our evaluation on ICU survivor samples and employed the mixed healthy HuS sample as the healthy control. We first examined whether ICU HuS collected from ICU survivors at admission confers quantifiable morphological changes to hMMTs. After a 6-day exposure to serum, hMMTs were fixed and stained with SAA and Hoechst 33342 for morphometric analysis at day 12 of differentiation (Figure 1A). This revealed the presence of multi-nucleated and aligned myotubes (Figure S3) and comparable nuclear fusion indices (Figure S4) across all samples tested.

**Figure 1.**
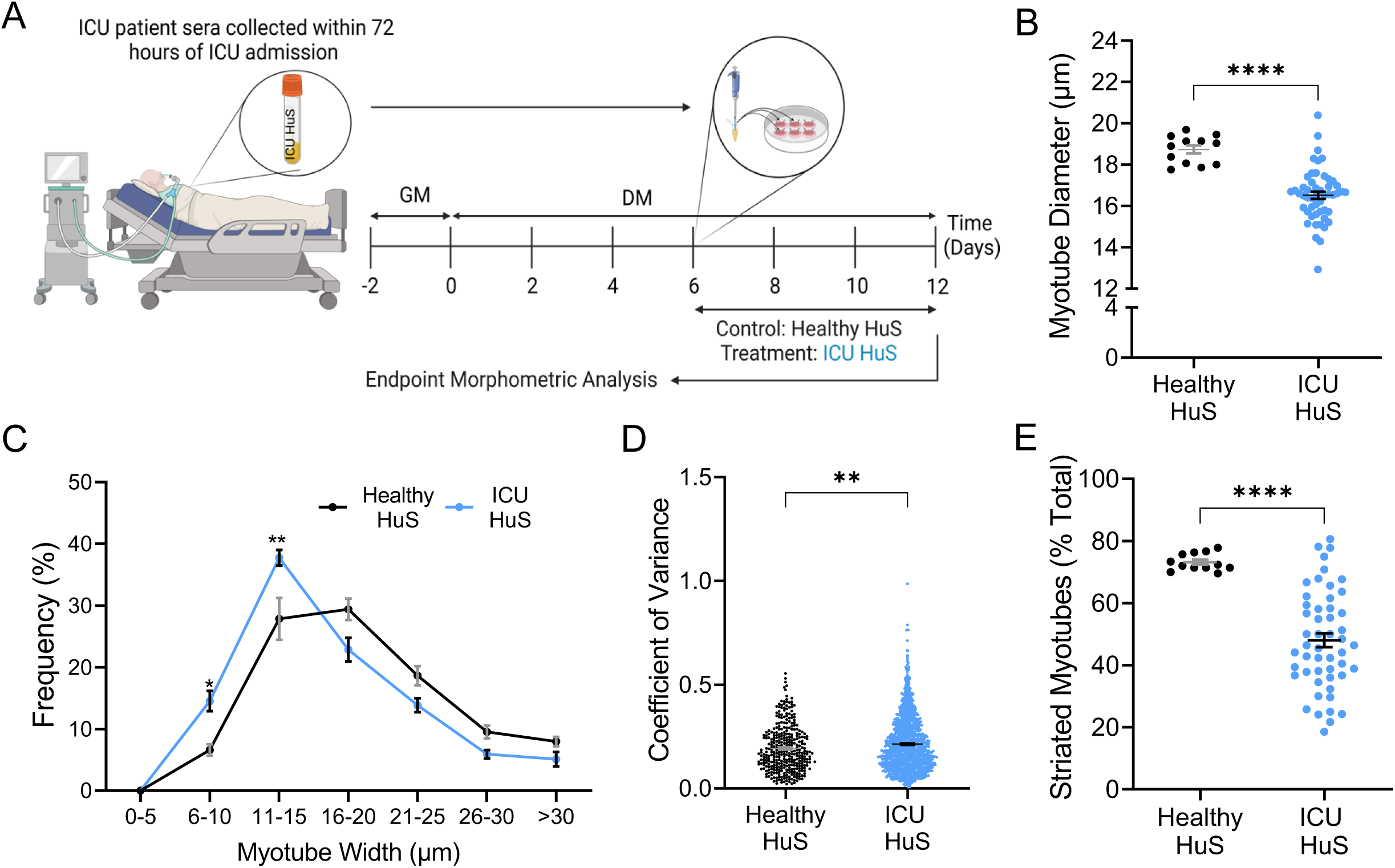
hMMT morphometric quality declines in response to ICU admission serum. A) Schematic overview of hMMT morphometric analysis timeline (made with BioRender). B) Dot plot quantification of myotube width analysis where each dot indicates the average myotube diameter quantified within single hMMTs. Each dot represents mean myotube width for an individual hMMT. C) Frequency polygon of myotube size distribution between Healthy HuS and ICU HuS treated hMMTs. Mean with SEM is shown. *** p < 0.001 and * p < 0.05 determined by 2way ANOVA using Tukey’s multiple comparisons test. D) Dot plot of myotube width variation indicated by the coefficient of variance. Each dot represents the coefficient of variance for a single myotube with an hMMT. n = 352 myotubes from Healthy HuS, n = 1508 myotubes from ICU HuS. E) Dot plot displaying the percentage of striated myotubes in hMMTs treated with healthy HuS or ICU HuS. Each dot represents mean % striation for an individual hMMT. B, E). n = 12 hMMTs treated with healthy HuS, and n = 51 hMMT treated with ICU HuS. B,D-E) N = 17 ICU patient sera samples. Mean with SEM is shown. **** p < 0.0001, ** p < 0.01, * p < 0.05, by unpaired student t-test.

Within days of ICU admission, muscle mass is reported to fall rapidly ^43^. We observed a significant decline in rectus femoris cross sectional area in our patient cohort when comparing ICU admission values with those at ICU discharge (Figure S5). We also observed a statistically significant reduction in the diameter of myotubes within the hMMTs treated with ICU HuS samples compared to those treated with healthy HuS (Figure 1B-C). Furthermore, the variance in myotube width increased significantly with ICU HuS treatment, signaling a potential decline in sarcomere organization and/or content (Figure 1D). Indeed, sarcomere disorganization has been reported at the single fiber level in muscle biopsies from ICUAW patients ^20,44,45^. Therefore, we next evaluated the percentage of striated myotubes in our ICU HuS treated hMMTs. We found that the proportion of striated myotubes was markedly lower in ICU HuS treated groups relative to healthy controls (Figure 1E). Collectively, this data demonstrates that hMMT treatment with ICU patient serum collected within 72 hours of admission induces severe atrophy and sarcomeric disorganization, features that are also associated with CIM.

### 4.3 hMMTs treated with admission serum from ICU survivors display decreased absolute force and impaired force kinetics

To investigate the influence of admission serum on hMMT contractile function, we conducted analyses of passive and induced force generation (Figure 2A). First, we calculated the passive force (i.e. hMMT intracellular tension) by comparing phase contrast images taken on day 6 (before HuS exposure) to those acquired on day 12, after the 6 days of HuS treatment. We found that hMMT intracellular tension declined from days 6 – 12 of differentiation in the presence of ICU HuS (Figure 2B).

**Figure 2.**
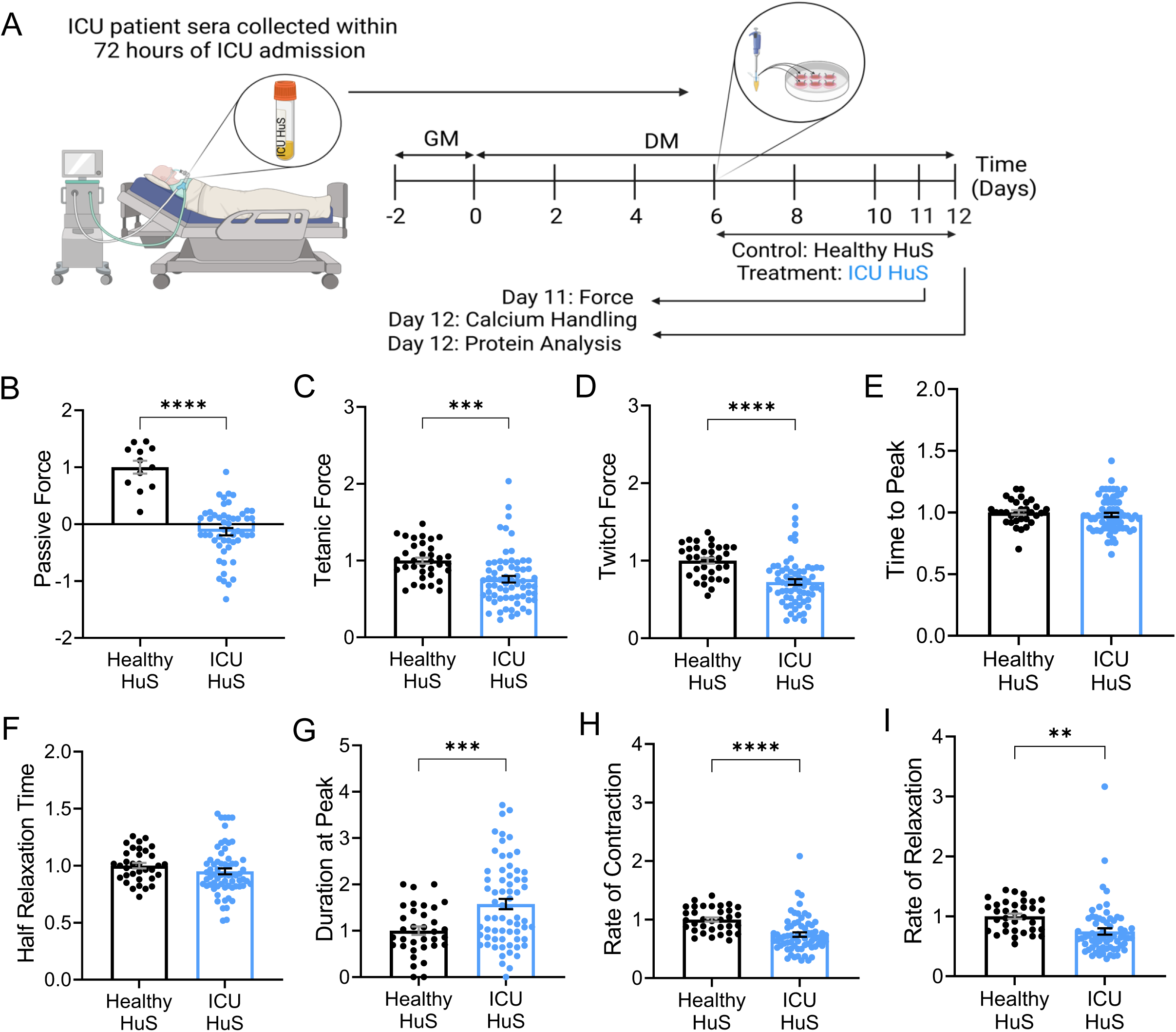
hMMT force generating capacity and contraction kinetics are diminished by treatment with ICU patient serum. A) Schematic overview of hMMT experimental timeline for Figures 1-3 (made with BioRender). B-I) Bar graph quantification of B) normalized passive force (n = 12 hMMTs for Healthy HuS, n = 51 hMMTs for ICU HuS with N = 17 ICU patient sera samples tested), C) normalized tetanic force, D) normalized twitch force, E) time to peak, F) half relaxation time, G) duration at peak, H) rate of contraction, and I) rate of relaxation determined on Day 11 of differentiation. Each dot represents mean data arising from a single hMMT. C-I) n = 34 hMMTs treated with Healthy HuS, n = 66 hMMTs treated with ICU HuS with N = 17 ICU patient sera samples tested. Mean with SEM is shown. **** p < 0.0001, *** p < 0.001, ** p < 0.01, * p < 0.05, by unpaired student test.

We next used our previously reported post-tracking script in Python to quantify and convert hMMT-induced post-deflection from videos captured during E-Stim on day 11 of differentiation into force values ^31,32^. Our studies revealed a significant decline in absolute force in response to high and low frequency E-Stim when hMMTs were exposed to ICU HuS (Figure 2C-D), and aligning with the diminished excitability reported for ICU patient muscle fibers ^17,18^. We then evaluated contraction kinetics at low frequency E-Stim. There was no discernable distinction across treatment groups in the time required for hMMTs to reach maximum force, nor in the time to relax halfway from their maximum contracted states (Figure 2E-F). However, a notable disparity was observed with regards to ICU HuS treated hMMT duration at peak, and in rates of contraction and relaxation when compared to the healthy controls (Figure 2G-I). This suggests that although hMMTs exposed to admission serum can achieve their maximum force and then relax halfway relatively quickly, the overall process of contraction and relaxation is slower compared to the healthy controls. This may be due to factors such as impaired calcium handling, reduced muscle fiber recruitment, or alterations in muscle fiber composition, each of which has been reported in CIM clinical studies and animal models ^18,28,46^.

### 4.4 Total calcium release is significantly reduced after treatment with admission serum from ICU survivors

Intracellular calcium handling impairments have been implicated in the pathophysiology of CIM^28,47,48^. Therefore, we next evaluated the calcium handling properties of hMMTs 12 hours after the conclusion of our force studies, coinciding with day 12 of differentiation (Figure 2A). hMMTs were loaded with a fluorescent and cell-permeable calcium indicator (Calbryte^TM^ 520 AM, 10 uM). hMMT intracellular calcium transients were recorded, and subsequently evaluated by normalizing the fluorescent intensity of each electrically stimulated tissue to its own baseline fluorescent intensity (ΔF/F_0_). Treatment with dantrolene sodium confirmed that the calcium indicator was measuring Ca^2+^ released from the sarcoplasmic reticulum of hMMT myotubes (Figure S6). Representative epi-fluorescent images captured during E-Stim, and line graphs illustrating twitch and tetanic calcium traces from a single healthy HuS hMMT are shown (Figure 3A-B,D). Peak ΔF/F_0_ induced by low and high frequency E-Stim trended toward a decline in hMMTs treated with ICU HuS with significance observed with high frequency E-Stim (Figure 3C,E). Irregular calcium transient traces were observed at high frequency E-Stim. Specifically, those treated with ICU HuS failed to sustain the tetanic stimulus, as indicated by a rapid decline in ΔF/F_0_ values followed by signal fluctuation (Figure 3F). To further investigate this phenomenon, we quantified the total Ca^2+^ released during a 2-second tetanic stimulus (Figure 3G). Our findings revealed a significantly lower total Ca^2+^ release during the 2-second tetanic stimulation in ICU HuS-treated hMMTs that was not observed for healthy control hMMTs (Figure 3F-G).

**Figure 3.**
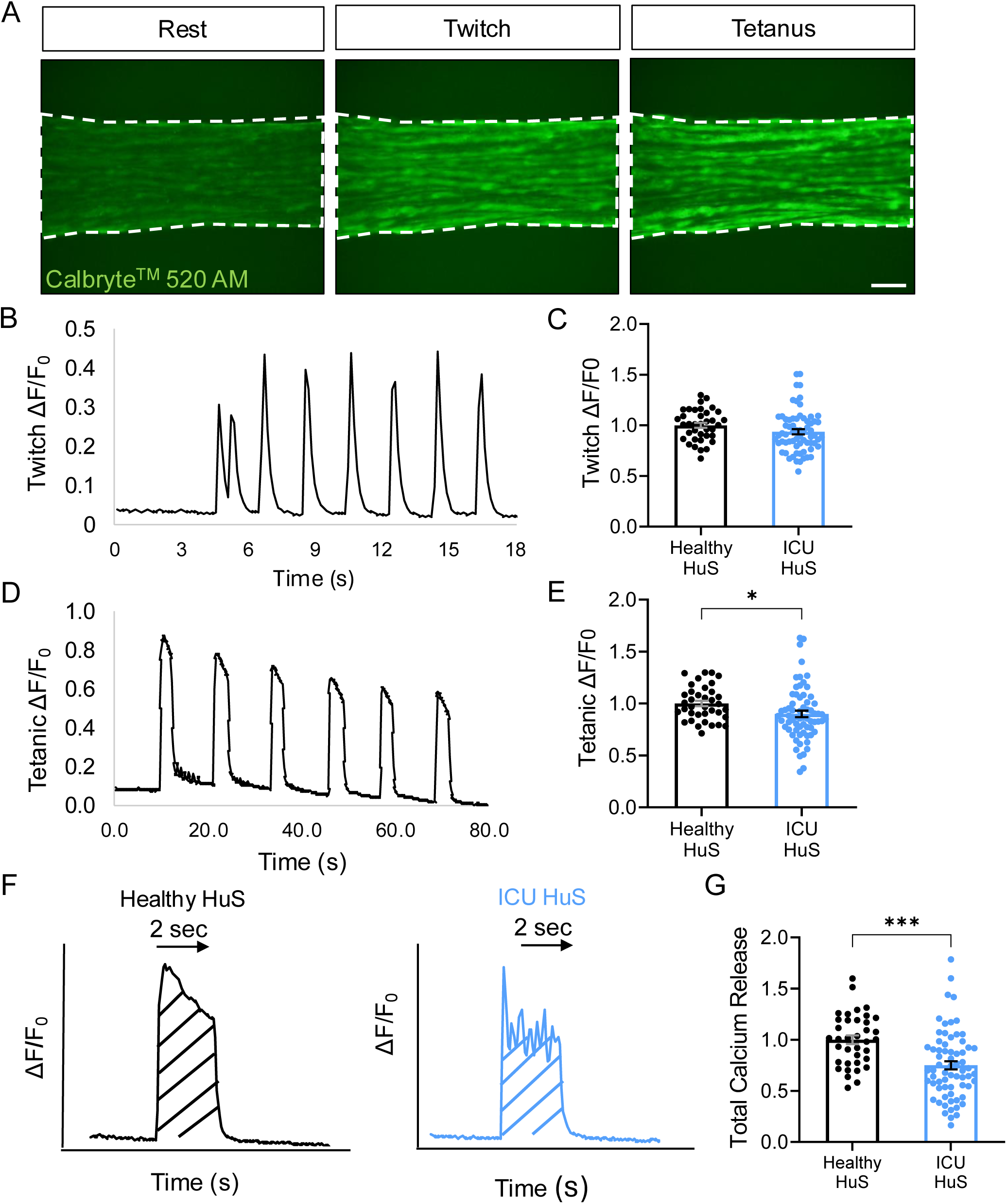
Total Ca^2+^ release during tetanic stimulation is significantly reduced in hMMTs treated with ICU patient sera. A) Representative epi-fluorescence images of peak Calbryte^TM^ 520 AM signals (green) at rest and during low (twitch) and high (tetanus) frequency electrical stimuli from Healthy HuS treated hMMTs. Scale bar, 50 μm. B, D) Line graphs showing representative intracellular calcium transient traces of Healthy HuS treated hMMTs in response to low (twitch, B) and high (tetanus, D) frequency electrical stimuli. C, E) Bar graph quantification of normalized peak intracellular calcium transients in response to low (C) and high frequency (E) electrical stimuli. F) Representative line graph of single tetanus calcium transient in Healthy HuS (left) and ICU HuS (right) treated hMMTs. Diagonal lines highlight the region used for integrated density analysis (mean grey value x area) over the 2-second tetanic stimulation period. G) Bar graph quantification of normalized integrated density (total calcium release) over a 2-second tetanic stimulation period. C,E,G) All graphs compare Healthy HuS and ICU HuS. Each dot represents a value quantified from a single hMMT. n = 37 hMMTs treated with Healthy HuS, n = 67 hMMTs arising from the analysis of N = 17 ICU HuS samples. Mean ± SEM is shown. *** p < 0.001, * p < 0.05, by unpaired student test.

Taken together, the force (Figure 2) and calcium handling data (Figure 3) presented herein suggest that admission serum induces hMMT functional deficits mirroring hallmarks of poor excitation-contraction coupling observed with CIM. These data indicate the suitability and potential of 3D muscle microtissues, over traditional 2D cultures, for modelling CIM *in vitro*.

### 4.5 Serum from a subset of ICU patients alters hMMT sarcomeric protein content

CIM has been associated with the preferential loss of thick filament protein myosin (MyHC), as well as reductions in the MyHC/Actin ratio ^5,17–19^. Therefore, we next investigated whether hMMT sarcomeric protein content was affected by treatment with serum from ICU survivors. Since we aimed to conduct protein studies using the tissues that underwent force and calcium transient analyses, we first confirmed that the E-Stim protocols did not alter MF protein content. Western blot analysis showed stable levels of pan myosin heavy chain (MyHC), SAA, and actin following two rounds of E-Stim (Figure S7). The subsequent analysis of sarcomeric proteins in hMMTs treated with ICU serum from survivors showed no statistically significant changes in pan-MyHC, SAA, actin, or in the MyHC/Actin ratio, relative to hMMTs treated with healthy control (Figures 4A-E). These data support our earlier assertion that reduced myotube width observed in admission serum treated hMMTs (Figure 1B-D) is driven by sarcomere disorganization (Figure 1E) rather than degradation (Figure 4) ^45^. However, it is important to highlight that while averaging the data revealed no statistically significant differences, a subset of individual patient samples drove significant reductions in sarcomeric protein content and MyHC/Actin ratios (Figure 4B-E).

**Figure 4.**
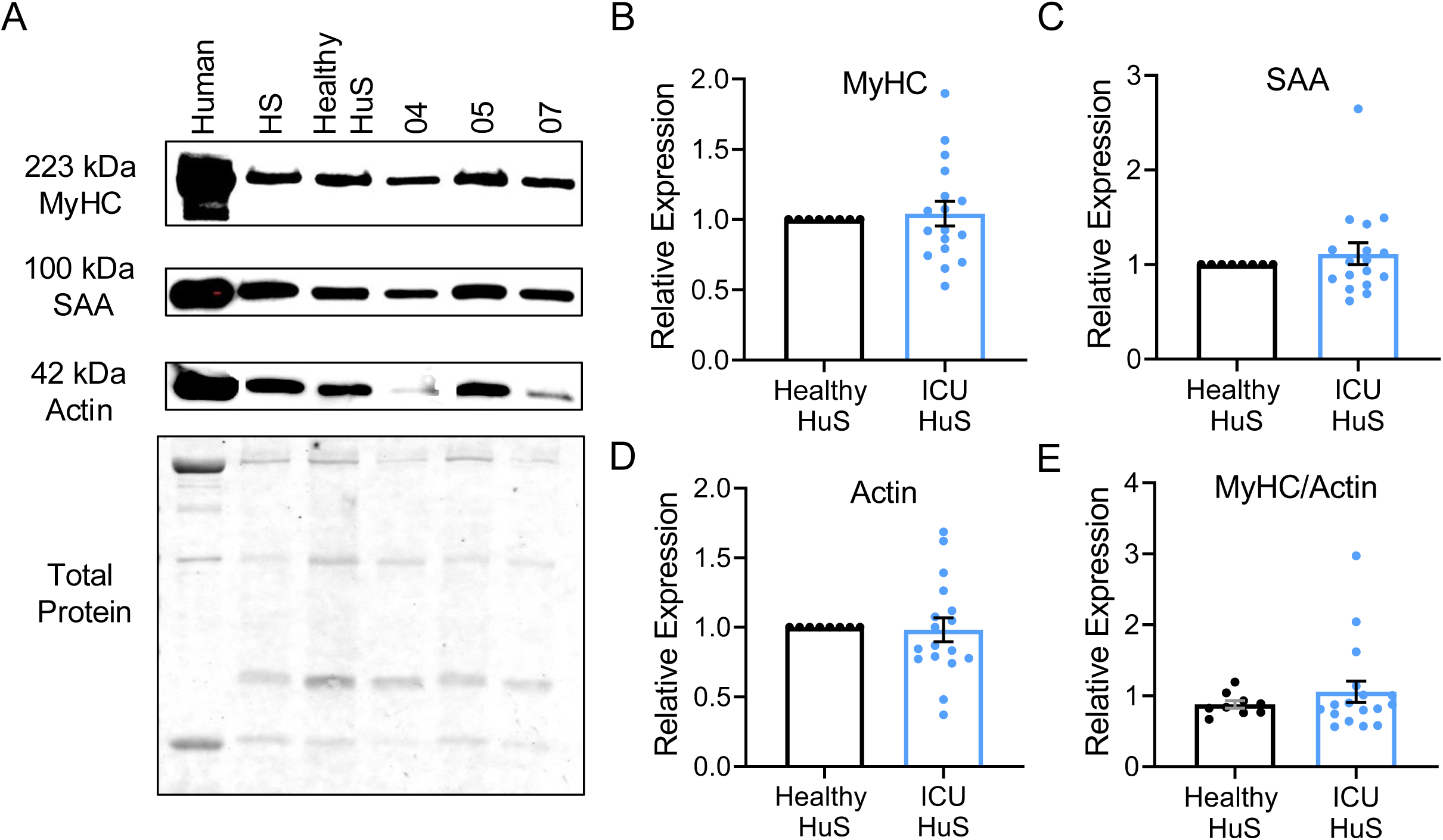
Mean myofibrillar protein content in hMMTs remains stable after ICU HuS treatment. A) Representative chemiluminescent blots (top) and ponceau total protein stain (bottom) from ICU patients 4, 5 and 7. B-D) Bar plots highlighting the relative B) myosin heavy chain (MyHC), C) sarcomeric α-actinin (SAA), and D) Actin expression in hMMTs normalized first to total protein and then to the healthy HuS control. E) Bar plot displaying the ratio of MyHC to Actin content in Healthy HuS and ICU HuS treated hMMTs. Values normalized to total protein. B-E) Mean ± SEM is shown. Lack of significance was determined by unpaired student t-test.

### 4.6 Individual patient level *in vitro* outcomes cluster with clinical risk factors

Our interrogation of hMMT morphological and functional responses to treatment with ICU patient admission serum indicates that humoral factors are capable of driving aspects of CIM pathogenesis. While statistically significant hMMT decline was clear from these population level studies, the patient-specific magnitude of effects varied. Therefore, we next aimed to explore these individual ICU patient patterns, and understand how they fit within the broader clinical context. For this, we unblinded our study to patient demographics and then related *in vitro* outcomes with well-known clinical risk factors such as age, ICU length of stay (LOS), and steroid treatment ^2,49–51^. We visualized this complex data using heatmaps (Figures 5 and 6), supplemented by bar plots to illustrate trends within the binned groups (Figure S8-9). A value of 1 indicates equivalency to the healthy HuS control and are coloured white. Values > 1 are orange (darker shades for higher values), and < 1 are blue (darker shades for lower values). We set thresholds at 1.5 (darkest orange) and 0.5 (darkest blue).

**Figure 5.**
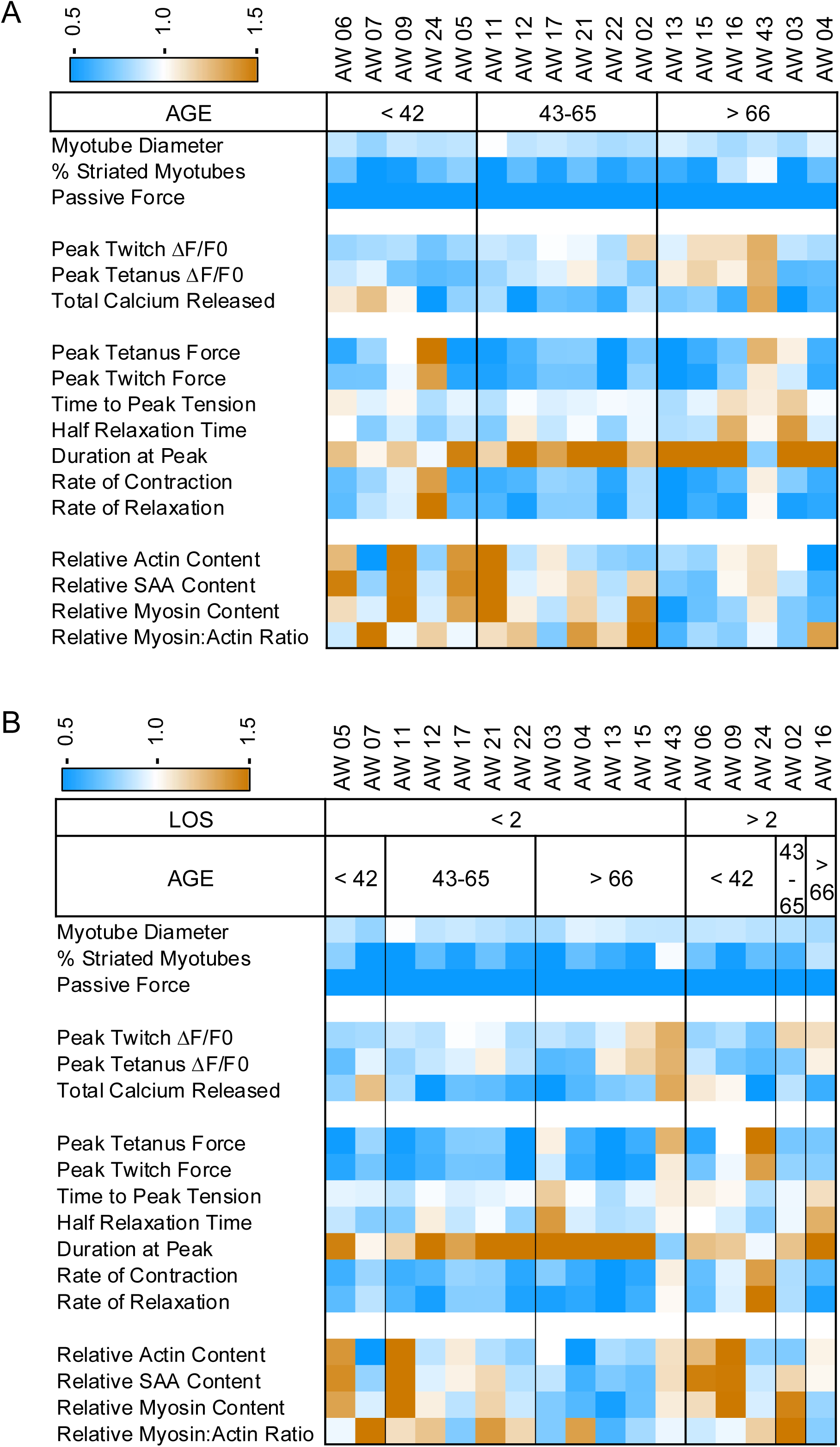
A distinct microphysiological signature distinguishes hMMTs treated with serum from ICU patients aged ≥ 66 who have been in the ICU for ≥ 2 weeks. A-B) Heats maps illustrating hMMT analysis data clustered by A) age or B) age plus length of stay (LOS). All values are normalized to their respective controls to ensure a consistent scale. A value of 1 indicates equivalency to the healthy human serum control and is white in colour. Values > 1 are orange with darker shades correspond to higher values). Values < 1 are blue with darker shades corresponding to lower values. Thresholds set at 1.5 and 0.5.

**Figure 6.**
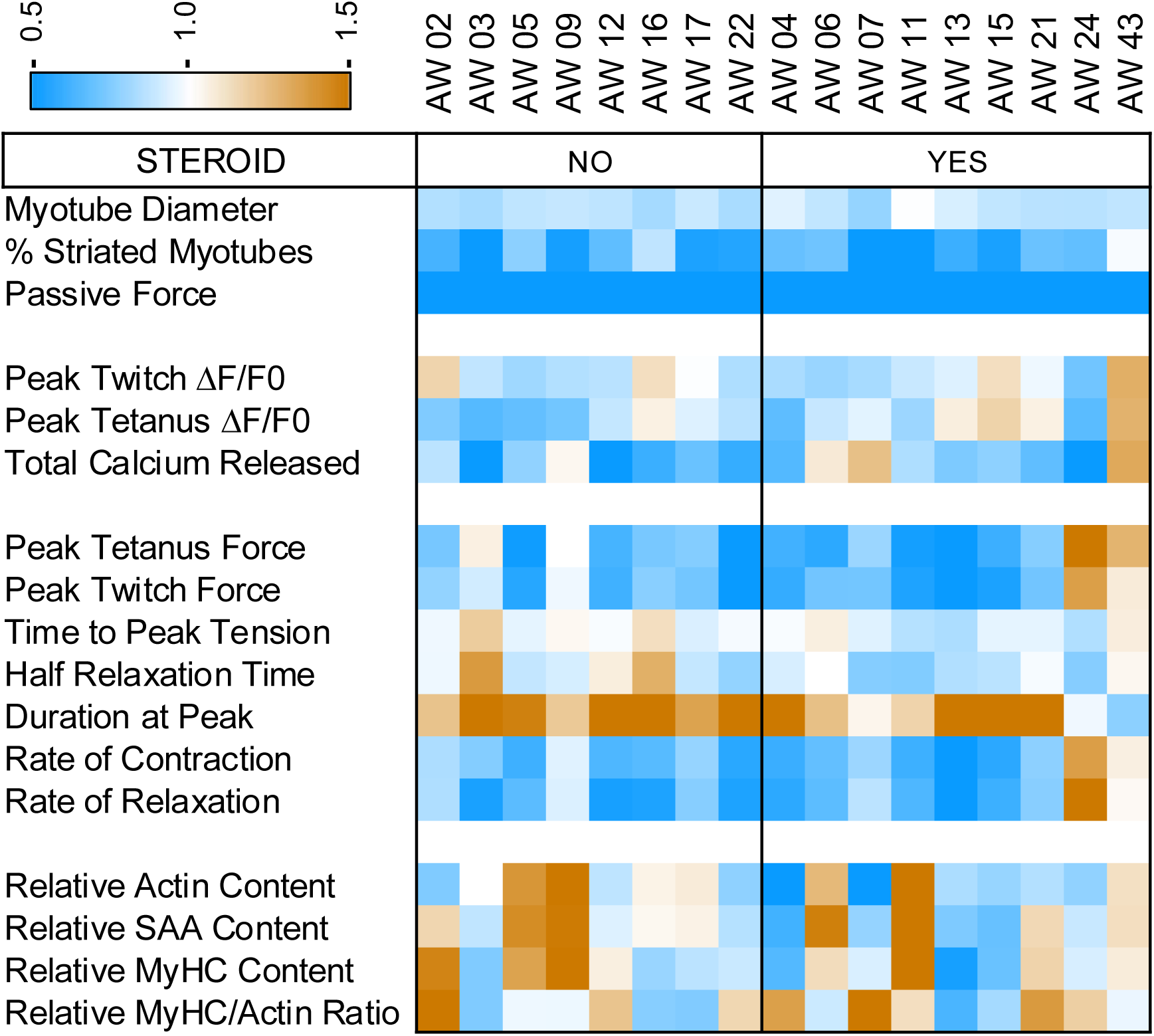
hMMTs exhibit impaired excitation-contraction coupling when treated with ICU HuS from those patients administered steroids in the ICU. Heats map illustrate hMMT analysis data clustered by steroid use. All values are normalized to their respective controls to ensure a consistent scale. A value of 1 indicates equivalency to the healthy human serum control and is white in colour. Values > 1 are orange with darker shades correspond to higher values). Values < 1 are blue with darker shades corresponding to lower values. Thresholds set at 1.5 and 0.5.

Age and ICU length of stay (LOS) strongly influence long-term patient functional outcomes ^13^. Previous research using the Functional Independence Measure (FIM) identified risk groups based on age and ICU LOS using recursive partitioning modeling^13^. Younger patients (< 42) with shorter ICU stays (< 2 weeks) had a higher chance of regaining full physical function, while older patients (≥ 66) with longer ICU stays (≥ 2 weeks) were less likely to regain independence ^13^. A proportion of our study population completed the FIM at 7 days post ICU discharge which revealed a decreasing motor sub score, indicative of impaired physical function, with older age, and longer duration of ICU stay (Figure S10). When clustered by age, loss of myotube diameter and striations were similar across all age groups (Figure 5A). Total calcium released by hMMTs was lower as the age of the ICU patient sera donor increased (≥ 42; Figure 5A, Figure S8A). A subtle loss of peak force (twitch and tetanus) was also observed (Figure 5A, Figure S8A) and these declines in absolute force were accompanied by impaired hMMT force kinetics conferred by sera samples from ICU patients with ages > 42 (Figure 5A, Figure S8A). Deficits in relative sarcomeric protein expression (actin, SAA, and MyHC), as well as the MyHC/Actin ratio, were increasingly apparent when ICU patient sera donor age was ≥ 66 (Figures 5A and S8A). The single outlier (AW 43; ≥ 66) was the only patient on mechanical ventilation for < 4 days.

We then extended our clustering analysis to include LOS as a parameter alongside age. In patients ≥ 66, MF protein loss occurred in response to admission sera regardless of the ICU patient sera donor LOS (< or ≥ 2 weeks; Figures 5B and S8B). Sera from older aged and longer LOS patients induced the greatest magnitude of hMMT force kinetic impairment (Figures 5B and S8B).

Lastly, we investigated associations when hMMT metrics were clustered by steroid use (Figure 6, Figure S9). The role of corticosteroid therapy in ICUAW remains controversial. Some clinical studies have indicated that corticosteroids may contribute to developing ICUAW^2,52,53^, yet others have demonstrated decreasing odds of developing ICUAW ^54^. While ICU HuS-treated hMMT myotube diameter and striations were consistent across patients, regardless of steroid treatment, several hMMT strength metrics trended higher for those who did not receive steroids in the ICU (Figures 6 and S9). The relative sarcomeric protein content of hMMTs treated with sera from ICU patients was not altered by steroid treatment, with the exception of MyHC, which trended higher in the absence of steroid treatment (Figures 6 and S9).

The outcome of this cluster analysis offers a convincing alignment between sera-induced *in vitro* phenotypes and two risk factors closely linked with recovery trajectories; age and length of ICU stay ^13^. By contrast, clustering based on steroid therapy revealed few associations suggesting that steroids and/or steroid induced changes to the humoral environment are not likely to be causing hMMT phenotypes. In sum, these observations serve to further strengthen the premise that humoral factors are likely to drive aspects of CIM-pathogenesis.

### 4.7 Admission serum from non-survivors significantly lowers the total calcium released by hMMTs while driving a mitochondrial response

We theorized that the admission serum obtained from ICU non-survivors (ICU-D) would impose distinct hMMT phenotypes compared to ICU survivor serum. Here we assessed this by focusing on the induced force and calcium handling properties of hMMTs treated with ICU-D serum, as well as to evaluate MF profiles. High frequency stimulations aimed at eliciting tetanic contractions revealed no notable differences when compared to healthy controls (Figure 7A). Low frequency contractions showed a trended decline in the peak twitch force of ICU-D HuS treated hMMTs compared to those treated with healthy HuS, but without reaching statistical significance (Figure 7B). Likewise, analysis of force kinetics during low frequency contractions revealed additional trended declines in time to reach maximum peak force (Figure 7C), half relaxation time (Figure 7D), duration at peak (Figure 7E), rate of contraction (Figure 7F), and rate of relaxation (Figure 7G) in the presence of ICU-D HuS. We then investigated the calcium handling activity of hMMTs treated with ICU-D admission serum. At low frequency E-Stim, the ICU-D serum treatment group showed a statistically significantly higher peak ΔF/F_0_ (Figure 7H). It is noteworthy that while both groups achieved comparable peak ΔF/F_0_ during high frequency stimulation (Figure 7I), the ICU-D serum treated hMMTs exhibited a significantly lower total Ca^2+^ release over a 2-second tetanic stimulation period (Figure 7J). Finally, we found that sarcomeric protein content remained stable in hMMTs after treatment with ICU-D HuS (Figure 7K-N).

**Figure 7.**
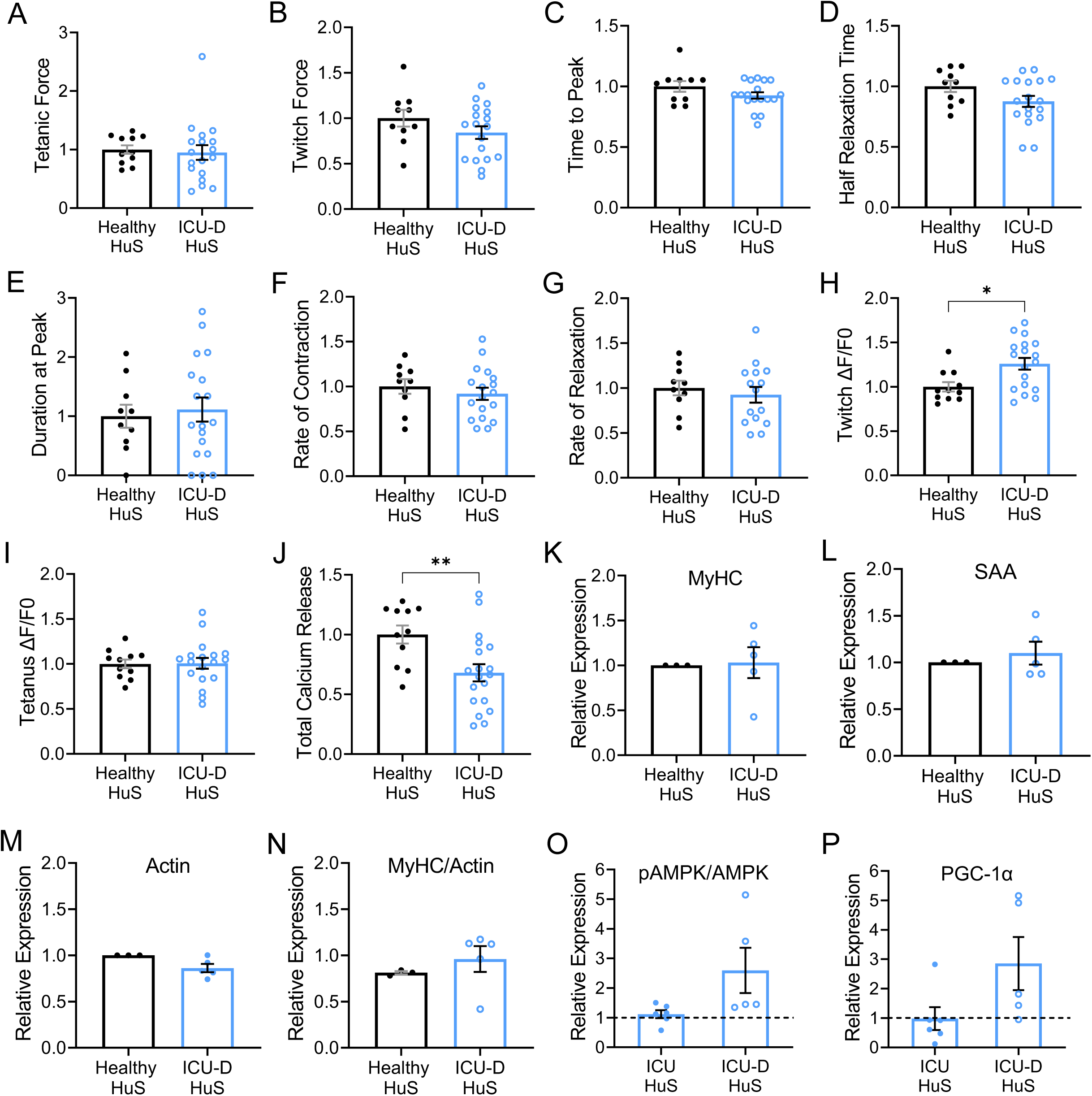
Serum from ICU non-survivors disrupts hMMT calcium handling and promotes a mitochondrial response. A-G) Bar graph plot depicting the quantification of individual hMMT force and force kinetics including normalized A) tetanic force, B) twitch force, C) time to peak tension, D) half relaxation time, E) duration at peak, F) rate of contraction, and G) rate of relaxation. H-J) Bar plots of hMMT calcium handling data visualizing normalized (H) peak twitch ΔF/F_0_, (I) tetanus ΔF/F_0_, and (J) total calcium release. A-J) Each dot represents a value quantified from a single hMMT. n = 10 hMMT treated with Healthy HuS and n = 18 hMMTs exposed to sera from N=5 ICU non-survivors (ICU-D) patient samples tested. K – P) Bar plots highlighting the relative K) myosin heavy chain (MyHC), L) sarcomeric α-actinin (SAA), M) Actin, N) ratio of MyHC to Actin, O) pAMPK/AMPK, or P) PGC-1α expression by hMMTs treated with Healthy HuS compared to those treated with ICU non-survivors (ICU-D). Values were normalized to total protein and then to the Healthy HuS samples (dotted line). Mean ± SEM is shown. ** p < 0.01, * p < 0.05 by unpaired t-test.

In conclusion, we report minimal alteration to hMMT function, or to MF protein levels, following treatment with admission serum from ICU non-survivors. This may reflect upregulated compensatory mechanisms. Indeed, upon evaluating signal transduction pathways associated with mitochondrial activity, we found upregulation of pAMPK/AMPK and PGC-1α protein expression by hMMTs treated with ICU serum from non-survivors, but not those treated with serum from ICU survivors (Figures 7O-P and S11). Interestingly, dysregulated pAMPK and PGC-1α levels was induced by serum collected at admission regardless of whether patient death occurred in under 10 days or after 25 days. These data suggest that the molecular response of hMMTs to admission serum collected from ICU patients may be used to augment APACHE II scores when advising family on ICU patient mortality rates.

## 5. Discussion

The specific role of circulating factors in ICUAW development remains poorly understood, and human microphysiological models that faithfully recapitulate this clinical disease are lacking. Here, we address both gaps using a novel 3D human skeletal muscle microtissue (hMMT) platform to dissect CIM biology driven by ICU patient sera. Remarkably, less than one drop of patient serum (8 µl total over 6 days) induced hallmark CIM features in hMMTs, demonstrating the direct pathogenic impact of humoral factors. Serum collected within 72 hours of ICU admission drove severe myotube atrophy and sarcomere disorganization, while functionally impairing Ca2+ release and force generation—effects observed in both survivors and non-survivors but with distinct severity patterns. Critically, we identified a functional and molecular compensatory signature that predicted patient mortality. hMMT *in vitro* metrics correlated strongly with established clinical risk factors including ICU length of stay, patient age, and steroid usage, validating the predictive power of our platform and revealing the potential to uncover new mechanistic insights into ICUAW pathophysiology.

The observed impairments in hMMT structural integrity and function following direct exposure to ICU patient serum provide compelling evidence for the involvement of blood borne factors in the patho-mechanism of CIM. The first evidence of a humorally mediated factor implicated in CIM was provided when researchers investigated the effects of blood serum fractions collected from five critically ill sepsis patients on the contractile functions of intact and skinned murine muscle fibers ^28^. Here they showed that pooled serum from CIM patients affected membrane excitability and the excitation-contraction coupling process at the level of the sarcoplasmic reticulum Ca^2+^ ^28^. Our calcium handling studies align with their observations, while delivering extended functional, morphometric, molecular, and sarcomere organization analyses. Moreover, by investigating the effects of each patient’s sera on hMMTs individually, we were able to discern patient-specific effects. While our investigation did not directly assess the effects of patient serum on muscle membrane excitability, we have established methodologies that pave the way for future exploration in this direction ^55^.

A distinguishing feature of muscle loss in CIM is the selective reduction of myosin and myosin-related proteins in comparison to actin, yet the underlying reasons for this phenomenon remain elusive. Many preclinical models of muscle atrophy induced by steroid exposure or muscle unloading fail to replicate this observation ^56^. Only within a single rodent model, where animals are subject to MV along with deep anesthesia and other common ICU stressors for extended periods ranging from days to weeks, has the preferential loss of myosin been documented ^25^. This finding strongly suggests that the replication of this phenomenon observed in humans requires a convergence of pathways, as well as a sufficient duration of exposure to these stressors. Despite six days of treatment with ICU serum, we did not observe a significant loss of MyHC or MyHC/Actin ratio at a population level in our advanced culture model. We did however, observe significant atrophy of hMMT, with reduction in diameter of myotubes treated with patient sera vs sera from healthy controls. Our observation that the rectus femoris cross sectional area declined in all patients in whom serial measurements in the ICU were obtained (Figure S5), further supports the premise that humoral factors induce a muscle response to critical illness.

Prior studies have reported that mechanical silencing is an important factor in triggering CIM ^57,58^. Mechanical silencing is characterized by the absence of external strain related to muscle loading, and internal strain generated by myosin-actin activation ^57,58^. In our culture model, we experience spontaneous hMMT contraction, possibly due to the immature ion conductance, leading to internal myosin-actin activation. This feature of the system may prevent a preferential myosin loss. On the other hand, the diaphragmatic muscle, also affected during CIM, is always passively active and does not experience preferential myosin loss^26^. Future experiments can implement drugs such as N-benzyl-p-toluene sulphonamide (BTS, a myosin – II inhibitor) ^59^, botulinum toxin (used to help treat dystonia and spasticity) ^60,61^, or dantrolene sodium (a direct-acting muscle relaxant blocking calcium release from the sarcoplasmic reticulum) ^62^ during the 6-day ICU serum treatment period to prevent internal activation of the hMMTs, and more accurately recapitulate mechanical silencing. However, a longer treatment period with ICU serum may be required, and a limitation of currently available protocols is that decline in hMMT integrity begins in the third week of culture. Others have shown that downregulation and loss of sarcomeric stabilizing protein titin occurs with CIM and may occur before preferential myosin loss ^63^. Thus, it may be worth investigating titin expression. Finally, researchers have shown that myosin post-translational modifications precede preferential myosin loss and fiber diameters remained unchanged during the first 5 days of ICU intervention, but significantly reduced at 10 days ^64^, further suggesting that we may require a longer treatment period in our hMMTs to observe this preferential myosin loss.

The robustness of our approach lies in its ability to replicate established clinical patterns. A significant proportion of ICU patients experience muscle mass loss during critical illness, with the majority regaining some muscle mass after discharge ^65^. Our analysis of myotube diameter data demonstrates that our hMMTs closely mirror this clinical phenomenon. Irrespective of physical functional capacity before entering the ICU, many patients will experience a decline in their physical function during ICU and hospital admission, with most patients experiencing incomplete recovery following discharge, with heterogeneity amongst patients^51,66^. Our *in vitro* dataset mirrors this variability at an individual patient level. Additionally, there is a pronounced decrease in force measures and sarcomeric protein content, particularly among individuals aged 66 and older, as well as those who endured an ICU stay of two weeks or longer, aligning closely with clinical observations ^13^. Finally, we observed that despite an increase in myotube diameter, there was not always a corresponding increase in hMMT strength, consistent with findings demonstrating that muscle mass reconstitution did not correlate with a resolution of muscle weakness ^20^.

The study is limited by the fact that we were unable to undertake serial measures of muscle strength in the ICU to determine the presence, or absence of ICUAW and thus have insufficient data to enable interrogation of the relationship between “in vivo”/patient strength versus the in vitro hMMT metrics. ICUAW can be diagnosed if MRCSS is less than 48 and there is no other directly identifiable reason for weakness, but the absence of serial testing in 5 individuals did not permit following the scores to ICU discharge, and no in-ICU measures were obtained in the remaining 17 patients. However, the serial ultrasound measures of RF CSA provided robust data showing significant and rapid muscle loss in the ICU in keeping with the reported literature, thereby validating the suitability of this patient cohort for study of the myotoxicity of critical illness. Similarly, the FIM motor subscores of this cohort demonstrated alignment with reported literature with older age and longer length of ICU stage negatively impacting post ICU physical functional outcomes.

To conclude, our study provides evidence supporting the involvement of humoral factors in the pathogenesis of CIM, emphasizing the potential of humoral mechanisms as therapeutic targets for CIM. By studying the role of humoral factors *ex vivo* in human cells within an advanced culture system, we successfully recapitulate both structural and functional hallmarks of CIM that were previously challenging to investigate *in vitro*. Future investigations should focus on identifying specific humoral factors dysregulated during CIM to understand whether these disease-associated humoral factors are indeed responsible for direct pathogenic effects. Nevertheless, we present a “CIM in a dish” model to study the biology of CIM; a scalable companion to patient-derived tissue and animal studies to elucidate mechanisms driving muscle atrophy and weakness in critically ill patients on the path to discovering targeted therapeutics.

## 6. Contributions

PMG and JB conceived of the project and designed experiments. HL designed and performed experiments, analyzed data, and prepared figures. All authors contributed to data interpretation. JB consented participants for the ICUAW study, collected clinical samples, and prepared figures. RAG consented participants for western blot control muscle lysate and collected a clinical sample. JC organized and transferred clinical samples to HL. SM and KW performed and analyzed patient ultrasound studies. PMG, JB, and SM supervised and funded the research. HL, PMG and JB wrote the manuscript. All authors edited and approved the manuscript. Schematics were prepared using BioRender.com.

## Supporting information

Lad et al Supplemental Figures

## 7. Acknowledgements

This work was supported by an Ontario Graduate Scholarship, NSERC CREATE TOeP Award, Cecil Yip Doctoral Research Award, Jennifer Dorrington Award, a Wildcat Graduate Scholarship, and Faculty of Applied Science and Engineering Student Endowment Fund (APSC GSEF) Award to HL; a University of Toronto EMH Seed Grant to JB and PMG; CIHR MOV-408234 to JB, CDS, MH, PG; and the Canada Research Chair in Endogenous Repair to PMG.

## 8. Supplemental Figure Captions

**Supplemental Figure 1. hMMT width and myotube diameter are unaffected by healthy HuS**. A) Timeline for immortalized myoblast-derived hMMT growth and maturation highlighting when healthy human serum is present. B) Representative phase contrast images of hMMTs at 4x magnification on day 6 and day 12 of differentiation. Scale bar = 500 µm. C) Line graph quantifying the remodeling of hMMTs as an assessment of the compaction of hMMT width during differentiation with HS (Day 4), when HuS 1 was added on Day 6 (indicated by vertical dotted line), and an additional time differentiation timepoint (Day 10) as well as the assay endpoint (Day 12). n = 3 hMMTs for each condition. All values are reported as mean with SEM. No significance was determined using two-way ANOVA followed by Dunnett’s multiple comparison test to compare HuS 1 to the control (2 % HS), at days 10 and 12 of differentiation. D) Representative tiled confocal images taken at 10x magnification of Day 12 hMMTs treated for 6 days with either control HS or healthy HuS 1. Sarcomeric α – actinin (SAA) is shown in magenta, and Hoechst 33342 nuclear stain is shown in yellow. Scale bar = 500 μm. E) Dot plot represents quantification of myotube width analysis where each dot indicates the average myotube diameter quantified of a single hMMT at day 12 of differentiation. SEM is shown. No significance by unpaired t-test to comparing the mean of 2 % HuS1 to the 2 % HS control. n = 3 hMMTs per condition arising from N = 1 independent experiment.

**Supplemental Figure 2. Healthy HuS serum samples produce morphologically and functionally comparable hMMTs.** A) Representative 40x confocal images of myotubes formed in hMMTs at Day 12 of differentiation. Sarcomeric α – actinin (SAA) is shown in magenta, and Hoechst 33342 nuclear stain is shown in yellow. Scale bar = 50 μm. B-C) Dot plots showing quantification of B) myotube diameter, and C) percentage of striated myotubes across serum treatments. n = 9 hMMTs per condition arising from N = 3 independent experiments. D) Schematic diagram showing top and side view illustration of passive force outcomes as interpreted by MyoTACTIC post displacement. Posts are shown in grey and the microtissue structure is pink. Blue arrows demonstrate the direction of MyoTACTIC post-movement relative to Day 6 (left) in situations where tension increases (middle) or decreases (right). E) Bar graph quantification of absolute passive force where values above zero indicate an increase in passive force, while values below zero indicate a decrease in absolute passive force. n = 9 hMMTs for each serum analyzed from N = 3 independent experiments. F-G) Bar graph quantification of F) absolute DOF and G) tetanus contractile force produced by hMMTs on Day 12 of differentiation following exposure to control serum. n = 6 hMMTs analyzed for each healthy serum across N = 2 independent experiments. B-C, E, F-G) Mean with SEM is shown. No significance as determined by unpaired one-way ANOVA with Dunnett’s multiple comparisons test relative to the HS control.

**Supplemental Figure 3. hMMTs maintain striated and multinucleated myotubes after 6 days of treatment with ICU patient serum.** Representative 40x confocal images of immortalized hMMTs on Day 12 of differentiation following 6 days of treatment with ICU patient serum (N=18) or Healthy HuS (mixed control). hMMTs immunostained to visualize sarcomeric α-actinin (SAA, magenta). Insets in the top right corner show a 2x zoom of selected regions to highlight representative striation patterns. Scale bar = 50 μm.

**Supplemental Figure 4. Healthy HuS and ICU HuS treated hMMTs have comparable nuclear fusion indices.** Dot plot of nuclear fusion index analysis of healthy and ICU HuS treated hMMTs. Each dot represents the average nuclear fusion index of an hMMT. n = 12 hMMTs from healthy HuS, and n = 51 for ICU HuS arising from N = 17 ICU patient samples. Mean with SEM is shown. No significance was determined by the unpaired student t-test.

**Supplemental Figure 5. Significant decrease in patient rectus femoris cross-sectional area during ICU stay.** Paired dot plot of patient rectus femoris cross-sectional area as determined by ultrasound is shown at the time of study consent (within 72 hours of ICU admission; left column) and close to ICU discharge (right column). 11 of 22 patients had both measurements acquired. Missing measurements due to patient death, instability, operator unavailability or patient refusal. All data points are shown. ** p<0.01, paired t-test.

**Supplemental Figure 6. Dantrolene sodium inhibits Ca^2+^ release from the sarcoplasmic reticulum of hMMTs.** A) Representative epi-fluorescence images of calcium transients (green) arising from rest/no E-Stim, peak E-Stim (tetanus, 60 Hz), or E-Stim (tetanus, 60 Hz) + 75 nM dantrolene sodium treated hMMTs. Scale bar = 50 μm. B) Line graphs showing representative intracellular calcium transient traces of hMMTs in response to high frequency stimulation (tetanus, 60 Hz), with and without dantrolene sodium.

**Supplemental Figure 7. hMMT myofibrillar protein expression with or without E-stim.** A) Ponceau (right), cropped chemiluminescent (left) and raw chemiluminescent blot (right) to visualize myosin heavy chain (MyHC), sarcomeric α-actinin (SAA) and actin expression, as well as protein loading (Ponceau), in hMMTs following no electrical stimulation (E-Stim) or a single (1x) or double (2x) round of E-stim. B) Bar graph plots of protein expression normalized to total protein.

**Supplemental Figure 8. hMMT data related to ICU patient age and length of stay.** A-B) Bar graphs of data acquired for each *in vitro* hMMT analysis metric binned by age A) (< 42, 42-65, or ≥ 66) at the time of ICU admission of the sera donor or by B) age at time of ICU admission and length of stay (LOS; < 2 weeks or ≥ 2 weeks) of the sera donor. N = 17 patient samples. All hMMT data values are normalized to the corresponding healthy HuS control. Plots display median normalized values. Dots correspond to individual patient data arising from hMMTs treated with their sera. The Kruskal-Wallis test was initially employed to evaluate overall differences among age groups, and age + LOS for each metric. Subsequently, pairwise comparisons were conducted using Tukey’s Honestly Significant Difference (HSD) test to identify specific group differences.

**Supplemental Figure 9. hMMT metrics related to ICU steroid treatment.** A) Bar graphs of data acquired for each *in vitro* hMMT analysis metric binned by steroid treatment in the ICU (yes or no). N = 17 patient samples evaluated. All hMMT data values are normalized to the corresponding healthy HuS control. Plots display median normalized values. Dots correspond to individual patient data arising from hMMTs treated with their sera. The Mann-Whitney U test was utilized for its robustness against non-normally distributed data and its ability to assess differences between independent groups.

**Supplemental Figure 10. FIM motor sub score at 7 days post-discharge shows expected decline associated with older age and longer ICU stay.** Dot plot of patient FIM motor sub scores acquired at 7 days post-discharge. 10 of 22 patients had measurements acquired. Missing measurements due to patient unavailability or refusal. All data points are shown.

**Supplemental Figure 11. pAMPK expression is dysregulated in hMMTs treated with sera from ICU non-survivors.** A) Raw chemiluminescent (top) and ponceau stained (bottom) blots arising from hMMTs treated with sera from ICU non-survivors. B-C) Bar plots of B) AMPK and C) pAMPK (C) expression in hMMTs treated with Healthy HuS (dotted black line), ICU survivor sera (ICU HuS) or ICU non-survivor sera (ICU-D HuS). Values normalized to total protein. Significance was tested by unpaired t-test.

## Notes

### Competing Interest Statement

The authors have declared no competing interest.

