## Supplementary material for "Clinical risk factors of intensive care unit acquired weakness predicted by human muscle microtissue response to humoral factors": Lad et al Supplemental Figures

### SUPPLEMENTAL FIGURE 1

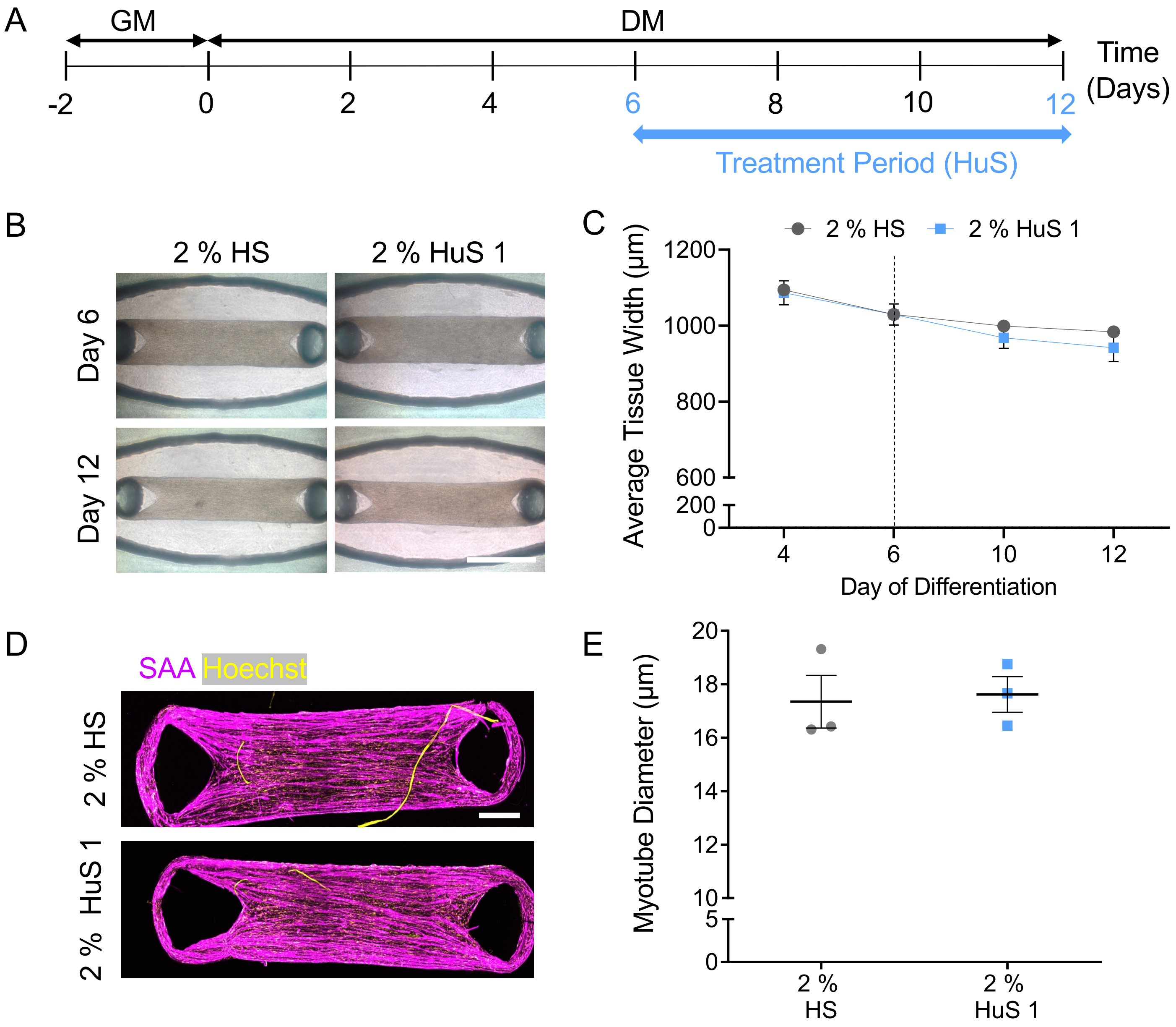

### SUPPLEMENTAL FIGURE 2

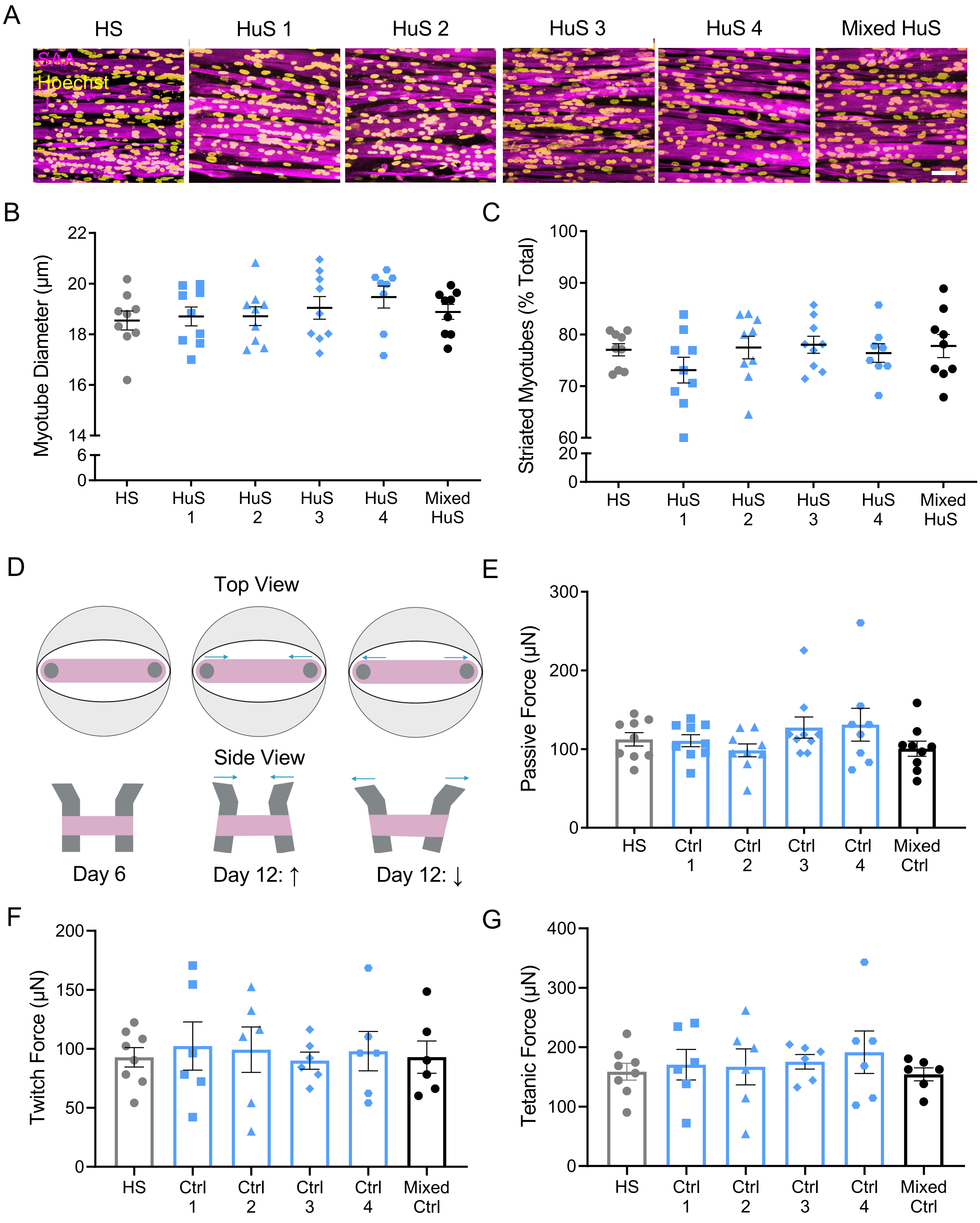

SUPPLEMENTAL FIGURE 3

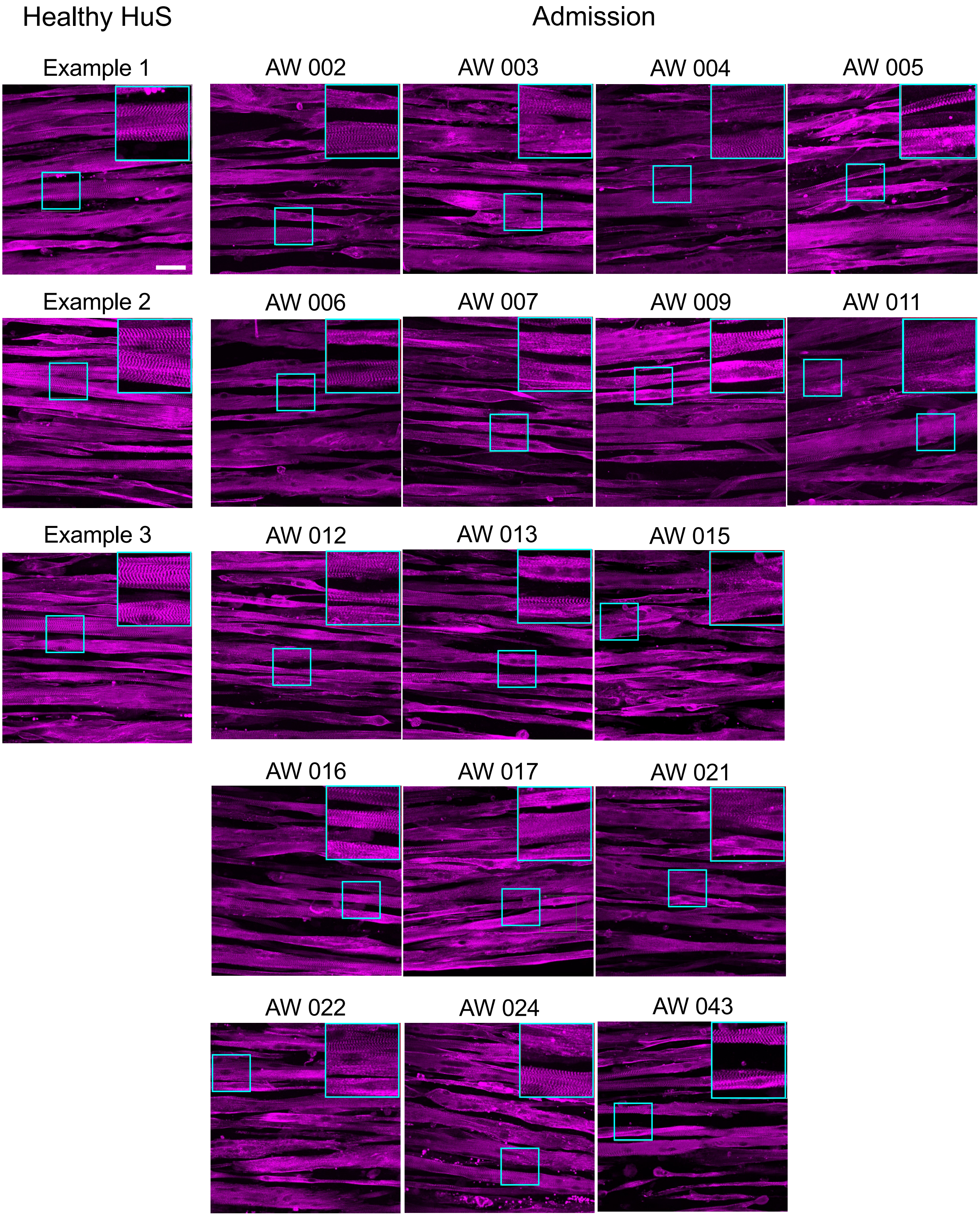

SUPPLEMENTAL FIGURE 4

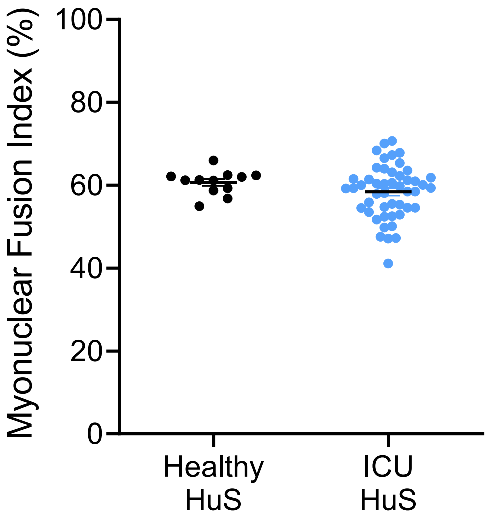

SUPPLEMENTAL FIGURE 5

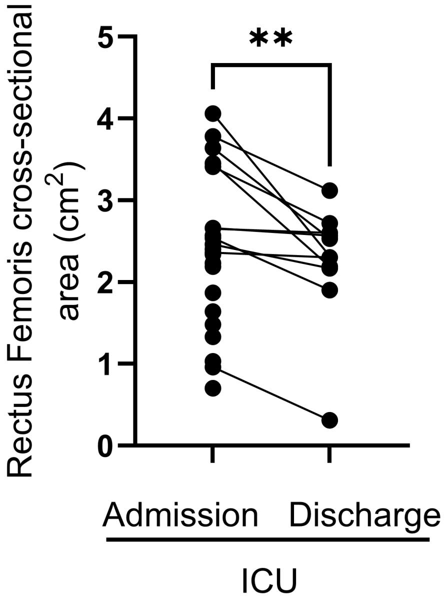

SUPPLEMENTAL FIGURE 6

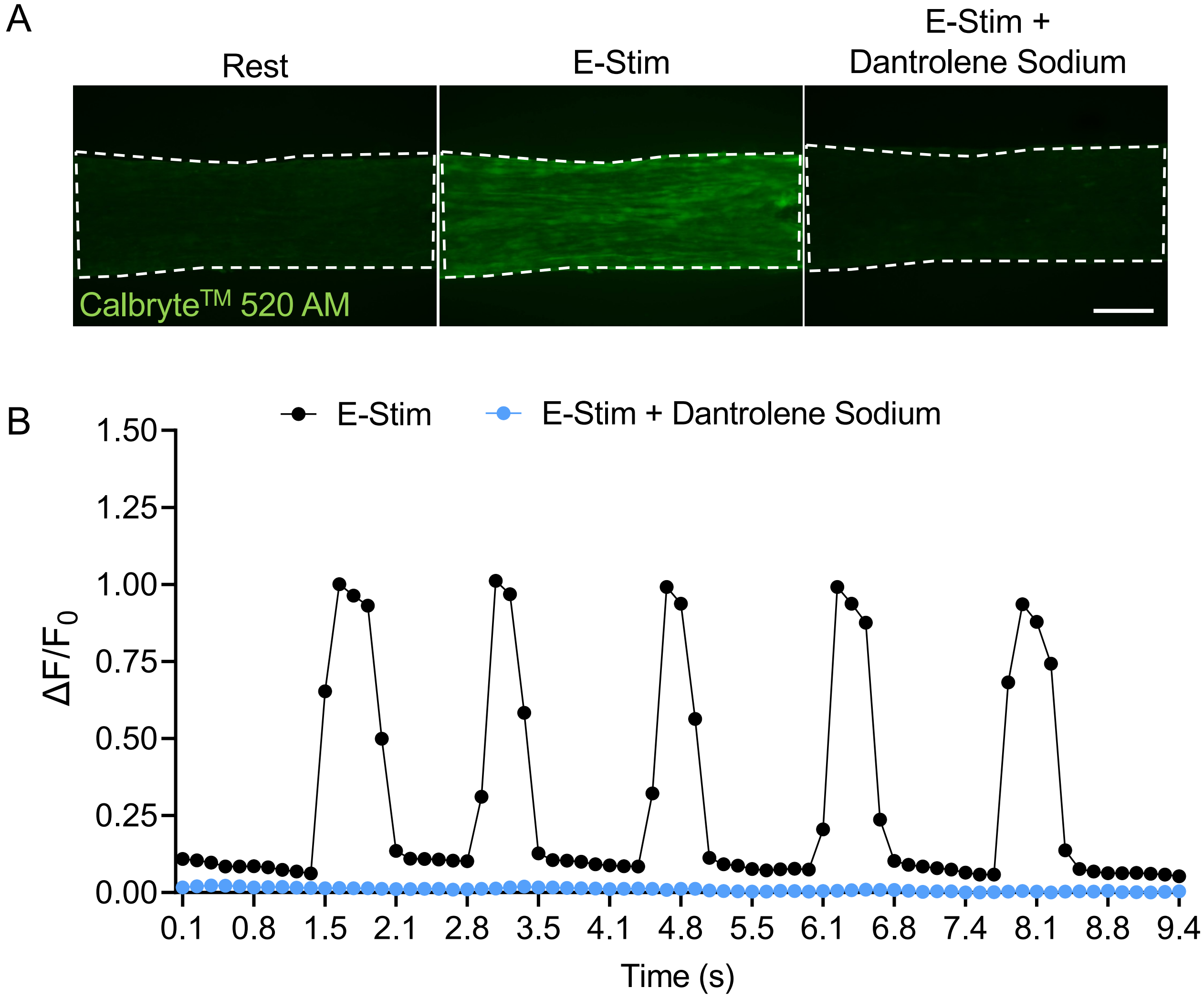

SUPPLEMENTAL FIGURE 7

A

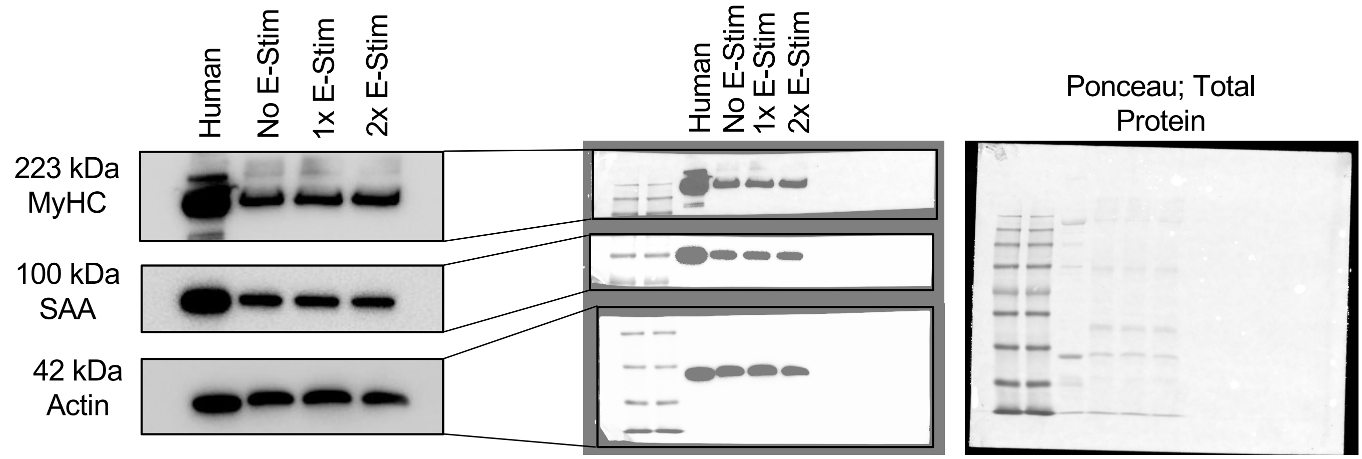


B

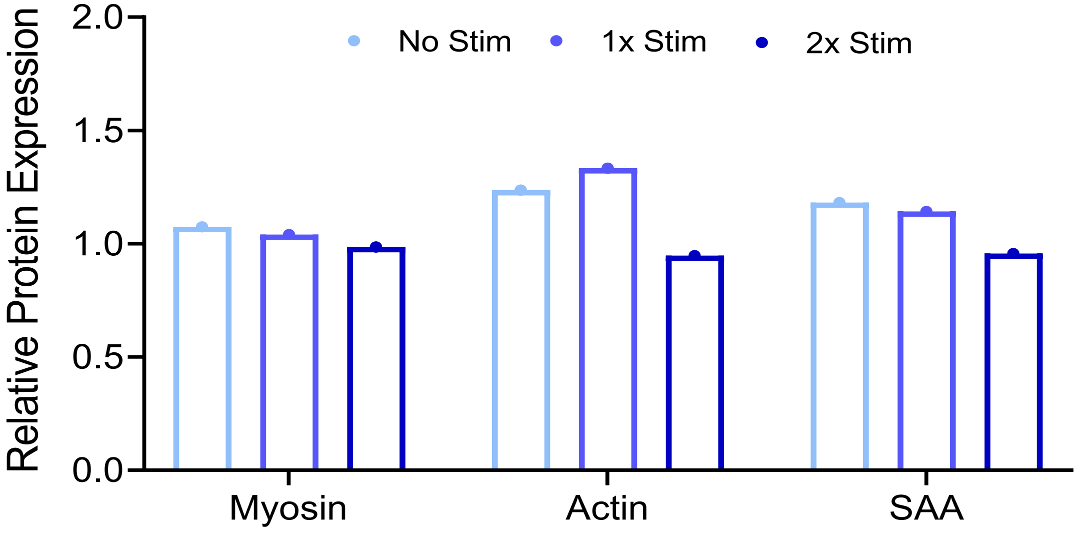

### SUPPLEMENTAL FIGURE 8

A

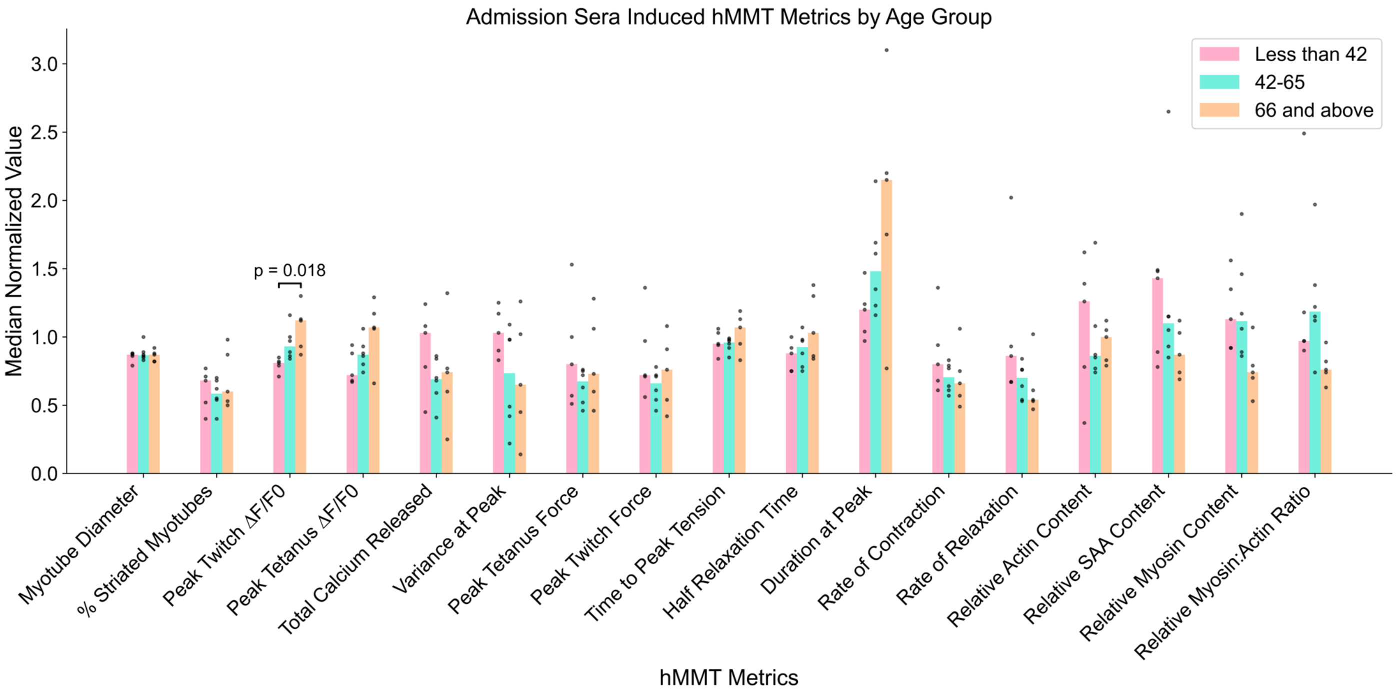


B

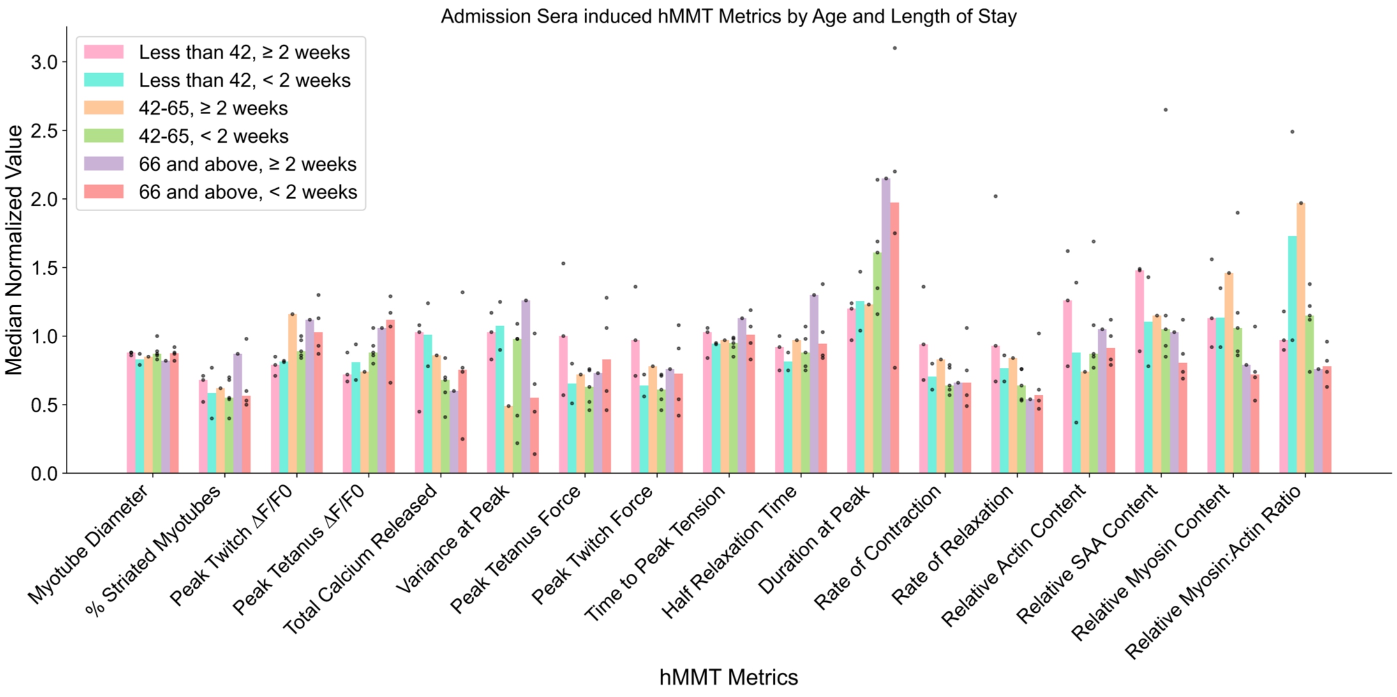

### SUPPLEMENTAL FIGURE 9

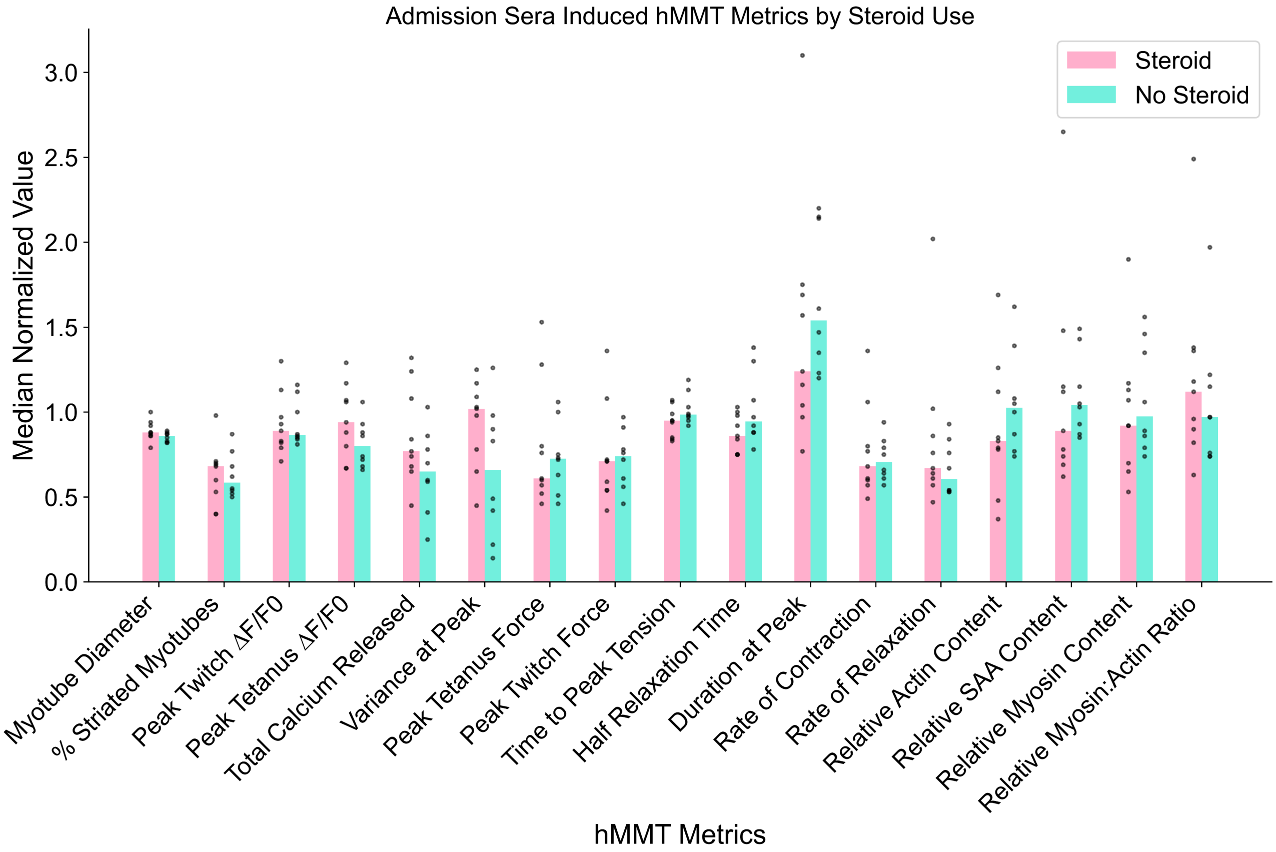

SUPPLEMENTAL FIGURE 10

A

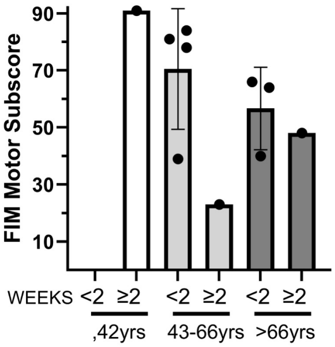

SUPPLEMENTAL FIGURE 11

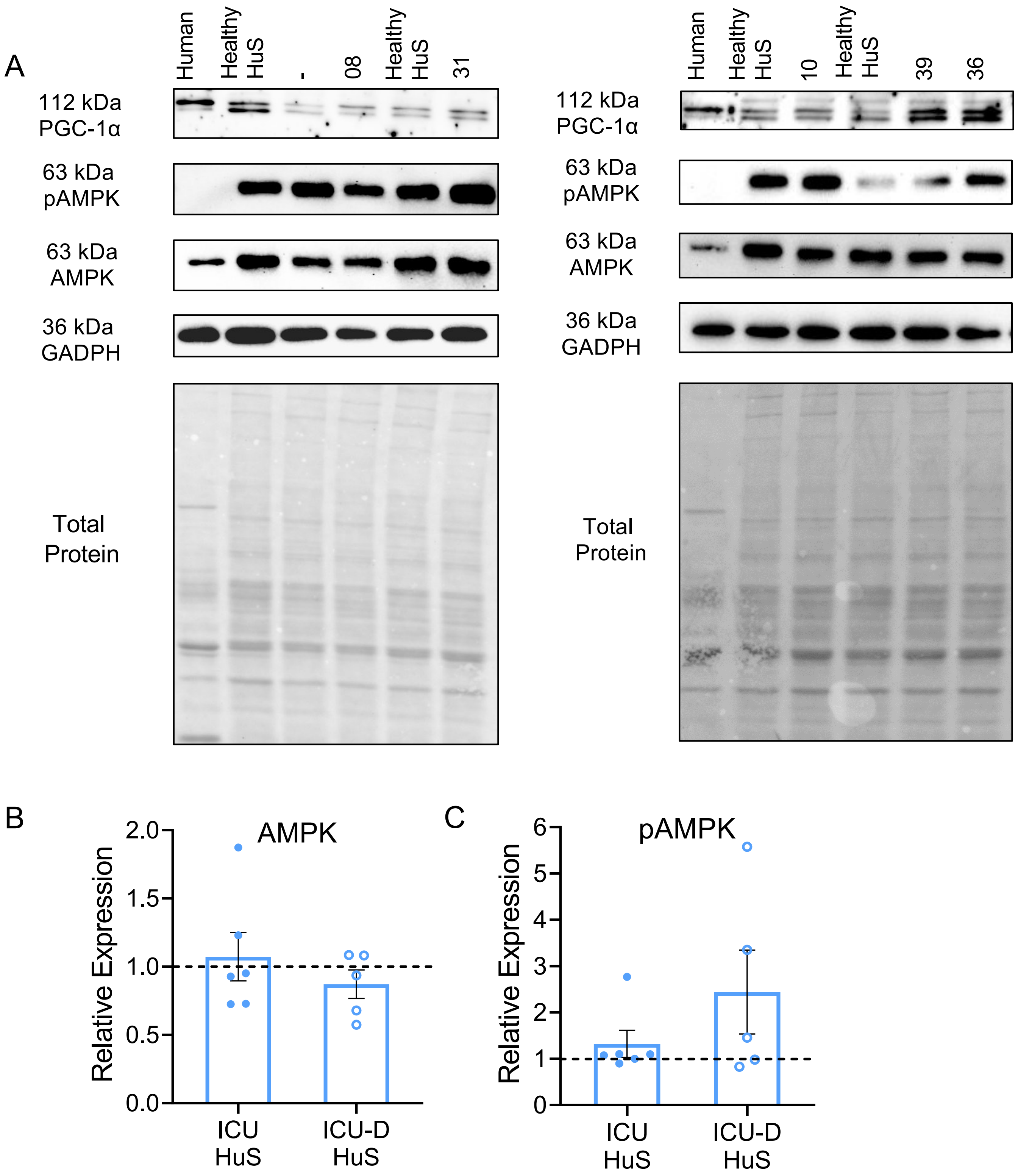
